# Nicotinamide mononucleotide (NMN) deamidation by host-microbiome interactions

**DOI:** 10.1101/2020.09.10.289561

**Authors:** Lynn-Jee Kim, Timothy J. Chalmers, Romanthi Madawala, Greg C. Smith, Catherine Li, Abhirup Das, Eric Wing Keung Poon, Jun Wang, Simon P. Tucker, David A. Sinclair, Lake-Ee Quek, Lindsay E. Wu

## Abstract

Oral administration of nicotinamide mononucleotide (NMN) is a prominent strategy to elevate nicotinamide adenine dinucleotide (NAD^+^) levels to treat age-related pathologies, where it is assumed to be directly incorporated into the NAD^+^ metabolome through the canonical recycling pathway. During oral delivery, NMN is exposed to the gut microbiome, which can modify the NAD^+^ metabolome through enzyme activities that are not present in mammals. Here, we show that orally delivered NMN can undergo direct deamidation and incorporation in mammalian tissue via the *de novo* pathway, and that this deamidation is reduced in animals treated with antibiotics to ablate the gut microbiome. Further, we show that antibiotics treatment increases the overall availability of NAD^+^ metabolites in the gut epithelium, with one possibility that the gut microbiome could be in competition with the host for dietary NAD^+^ precursors. Together, these data highlight previously undescribed interactions between orally delivered NMN and the gut microbiome.

## INTRODUCTION

Nicotinamide adenine dinucleotide (NAD^+^) is an essential redox cofactor central to metabolic processes such as glycolysis, the tricarboxylic (TCA) cycle and fatty acid oxidation^1, 2^. NAD^+^ is also consumed by enzymes such as the sirtuins^3^ and poly(ADP-ribose) polymerase (PARP) enzymes^4^ which are mediators of genome stability^5^ and DNA repair^6^. Given the essential role of this metabolite, the decline in NAD^+^ that occurs during biological ageing^7–12^ and disease states^13–15^ has gained attention as a target for therapeutic intervention^16, 17^. Strategies to boost NAD^+^ levels through supplementation with NAD^+^ precursors such as nicotinamide mononucleotide (NMN) and nicotinamide riboside (NR) are emerging as promising therapeutics^12, 16, 18–24^. Historically, dietary supplementation with the NAD precursors nicotinic acid (Na) or nicotinamide (Nam) was used to prevent chronic NAD deficiency, which causes pellagra. When these micronutrients are replete, the step converting Nam into NMN by the enzyme nicotinamide phosphoribosyltransferase (NAMPT) is thought to be rate limiting in NAD synthesis^25^, and the use of NAD precursors that occur after this step, namely NMN and NR, have gained prominence as a strategy to raise NAD^+^.

A surprising observation of oral delivery with the amidated metabolite NR is the sharp increase in levels of the deamidated metabolite nicotinic acid adenine dinucleotide (NaAD)^26^. NR is phosphorylated into NMN by NR kinases (NRK1/2)^27, 28^, and then adenylated into NAD^+^ by NMNAT enzymes (NMNAT1-3)^29–34^, effectively bypassing NaAD, which is an intermediate of the Preiss-Handler and *de novo* pathways^35, 36^. There is evidence from over half a century ago for the existence of enzymes in mammals that are capable of deamidating nicotinamide^37–39^ and NMN^38^, however the identity of these enzymes have not been revealed. In contrast, bacteria have a well-characterised NMN de-amidase enzyme, PncC^40^ that prevents the accumulation of NMN, which can be toxic to bacteria through its inhibition of the bacterial DNA ligase^41–43^. One explanation for the increase in NaAD with NR treatment^26^ could be that orally administered NMN and NR could undergo deamidation in the gut microbiome, yielding NaMN or NaR prior to their uptake into mammalian tissue, and then assimilation into NAD^+^ via an NaAD intermediate. A similar mechanism has been established for the administration of orally administered nicotinamide (Nam), whereby bacteria in the gut microbiome provide an alternate route to NAD biosynthesis through the deamidation of Nam into nicotinic acid (NA) by the enzyme PncA^44^, and there is also recent evidence for the ongoing exchange of NAD precursors between the gut microbiome and host tissues^45^. Notably, this exchange of NAD^+^ metabolites between the gut microbiome and the host has been implicated in the metabolism of orally administered NR^45^. A deamidation route mediated in part by the gut microbiome could explain the appearance of NaAD following NR supplementation^26^, however another unexplained aspect is that the delivery of labelled NR also results in the formation of NaAD^26^. This could suggest multiple mechanisms at play, including both direct deamidation of NAD precursors, and altered flux of NAD biosynthetic pathways following a bolus of exogenous NAD precursors.

Here, we used targeted metabolomics to trace the *in vitro* and *in vivo* metabolism of strategically designed NMN isotopologues to answer these questions. Consistent with recent work suggesting involvement of the gut microbiome in NAD metabolism^44, 45^, we show that NMN can be incorporated following its deamidation and metabolism via the *de novo* route, including both poorly characterised endogenous pathways and through mechanisms mediated in part by the microbiome. We further show that ablation of the microbiome by antibiotic treatment increases the uptake and conversion of orally delivered NMN into the NAD metabolome, and that isotope labelled NMN overwhelmingly presents in intestinal tissue in the form of NR. Contrary to the assumption that exogenous NMN treatment raises NAD^+^ levels solely through its direct incorporation into the NAD metabolome, we show that treatment with isotope labelled NMN increases the levels of endogenous, unlabelled NAD metabolites. Overall, our results provide unique insights into the assimilation of orally delivered, exogenous NMN into gastrointestinal tissue, and raise questions around how exogenous precursors alter the NAD metabolome.

## RESULTS

### NMN treatment alters the *de novo* arm of NAD^+^ synthesis

According to canonical models of mammalian NAD homeostasis, the metabolism of the amidated metabolite NMN does not intersect with the *de novo* pathway, which utilises deamidated intermediates (Fig. 1a). Unlike mammals, where putative nicotinamide (Nam) and NMN deamidase activity^37–39^ are poorly defined, bacteria present in the gut microbiome encode well-characterised deamidase enzymes such as PncC, which deamidates NMN into nicotinic acid mononucleotide (NaMN) for metabolism via the *de novo* pathway^40^. To test whether the gut microbiome alters the *in vivo* metabolism of orally administered NMN, we used mice that were exposed to a course of antibiotics to ablate the gut microbiome (Fig. 1b, Supp. Fig. 1). These animals received a bolus of NMN (500 mg/kg) by oral gavage, and four hours later, were sacrificed and tissues rapidly preserved for targeted metabolomic analysis (Fig. 1). We focused our analyses on the gastrointestinal tract (GIT) and the liver, as these two tissues have high levels of NAD synthetase (NADS) activity^46^, and are the primary sites of uptake and metabolism for orally delivered compounds. In agreement with previous work^26^, NMN treatment increased the de-amidated metabolites nicotinic acid riboside (NaR) and NaMN in both the gastrointestinal tract (GIT) (Fig. 1c, d) and liver (Fig. 1f, g), with an increase in nicotinic acid adenine dinucleotide (NaAD) in the liver only (Fig. 1h), matching previous findings for NR^26^. This spike in these deamidated metabolites was completely abolished in antibiotic treated animals, where NMN treatment instead led to a spike in the amidated metabolites NR (Fig. 1i, l) and NMN (Fig. 1j, m), and abolished the increase in liver NaAD (Fig. 1h). The abundance of each deamidated metabolite was expressed as a ratio of its amidated counterpart (Fig. 1o-t), highlighting an inverse relationship between amidated and deamidated metabolites during antibiotics treatment. Together, these data suggest a role for the microbiome in determining whether orally administered NMN is metabolised via the de-amidated or amidated arms of NAD metabolism.

**Figure 1.**
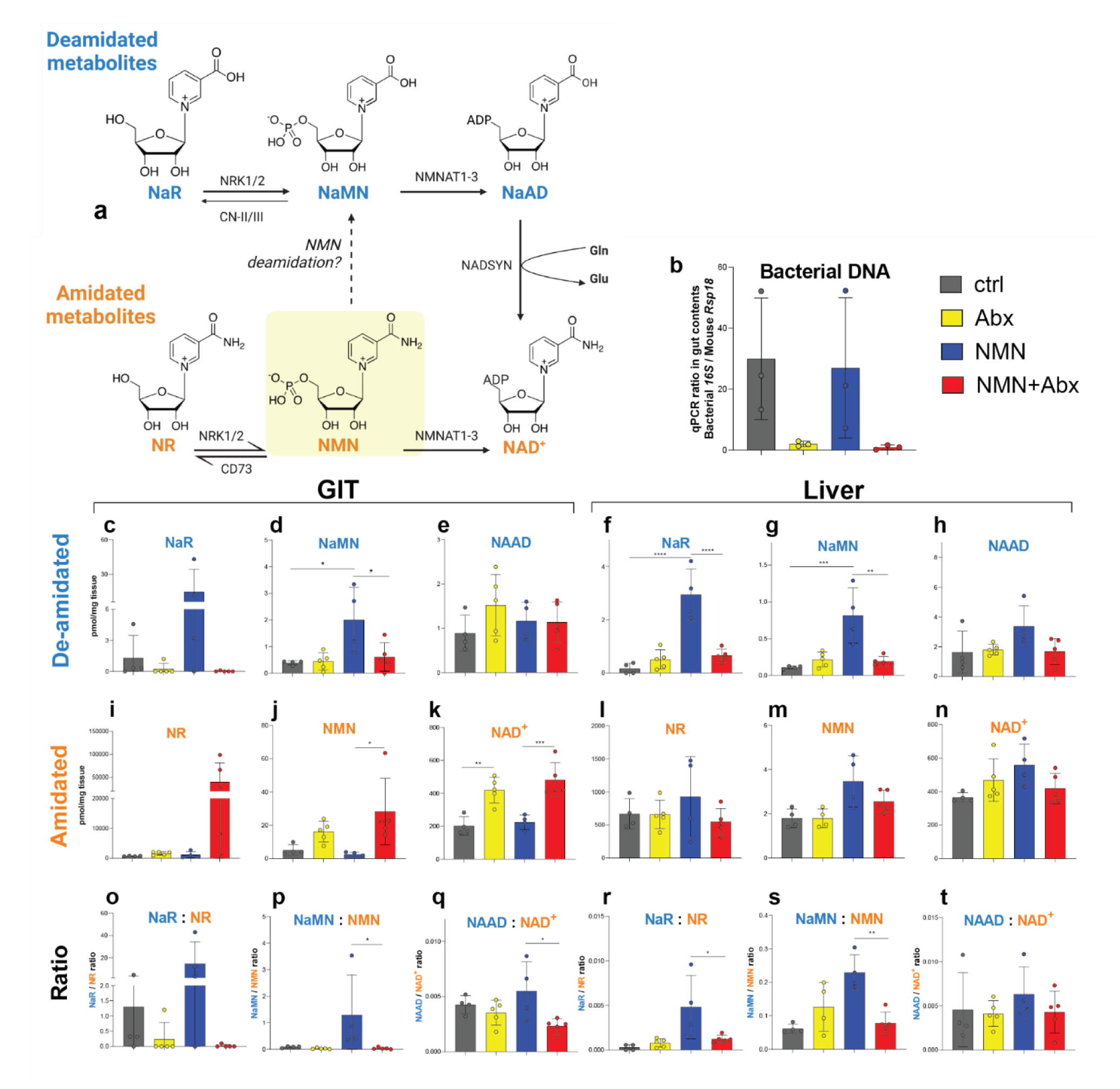
NMN treatment leads to the formation of deamidated NAD^+^ metabolites *in vivo* and is abolished by antibiotics treatment. (a) NMN is incorporated into NAD^+^ via the canonical recycling pathway, which does not involve deamidated metabolites present on the Preiss-Handler / *de novo* pathways. Mice treated with antibiotics (Abx) to ablate the gut microbiome, as determined by (b) qPCR for bacterial 16S DNA, and were administered a single dose of unlabelled nicotinamide mononucleotide (NMN) (500 mg/kg, oral gavage). Four hours later, (c-t) gastrointestinal tissue (GIT) and liver tissue were collected and subject to targeted mass spectrometry to quantify the deamidated metabolites (c-h) nicotinic acid riboside (NaR) (c, f), nicotinic acid mononucleotide (NaMN) (d, g) and nicotinic acid adenine dinucleotide (NAAD) (e, h), as well as their amidated counterparts (i-n) nicotinamide riboside (NR) (i, l), NMN (j, m) and nicotinamide adenine dinucleotide (NAD^+^) (k, n). These data were then expressed as ratios between de-amidated and amidated counterparts in GIT (o- q) and liver (r-t). Data analysed by 2-way ANOVA with Sidak’s post-hoc test, exact p-values and F values in supplementary files. N=4-5 animals per group, *p<0.05, **p<0.01, ***p<0.001, ****p<0.0001.

### Isotope tracing of NMN metabolism to reflect incorporation into NAD^+^ following deamidation

Given the impact of orally administered NMN on levels of deamidated NAD metabolites, we next sought to test whether this was due to the direct deamidation of NMN during its incorporation into the NAD metabolome, or was instead due to displacement of the deamidated pathway by an influx of an amidated precursor. To address this, we therefore conceived of three orthogonal approaches based on tracing the incorporation of isotope labelled metabolites for measuring the relative contribution of the amidated versus de-amidated pathways towards NAD^+^ biosynthesis, and the specific contributions of these pathways to the metabolism of exogenous NMN.

In the first strategy (Fig. 2a), we took advantage of the properties of the *de novo* NAD^+^ biosynthetic enzyme NAD synthetase (NADS), which amidates NaAD into NAD^+^ by replacing an O atom on the carboxylic acid of NaAD with an N atom from an ammonia intermediate derived from the amide group of the amino acid glutamine. Isotopic labelling of glutamine (Gln) with ^15^N at this amine group should result in the incorporation of this ^15^N label onto the amine of the nicotinamide ring of NAD^+^, resulting in an overall mass shift of M+1. In contrast, NAD^+^ biosynthesis via the recycling pathway does not utilise NADS, as the last metabolite in the pathway (NMN) is already amidated at this position. Depending on how much of the endogenous Gln pool is displaced with exogenous ^15^N-Gln label, the M+1 labelling of NAD^+^ at this position should therefore reflect the proportion of the NAD^+^ pool that was obtained via the enzyme NADS.

**Figure 2.**
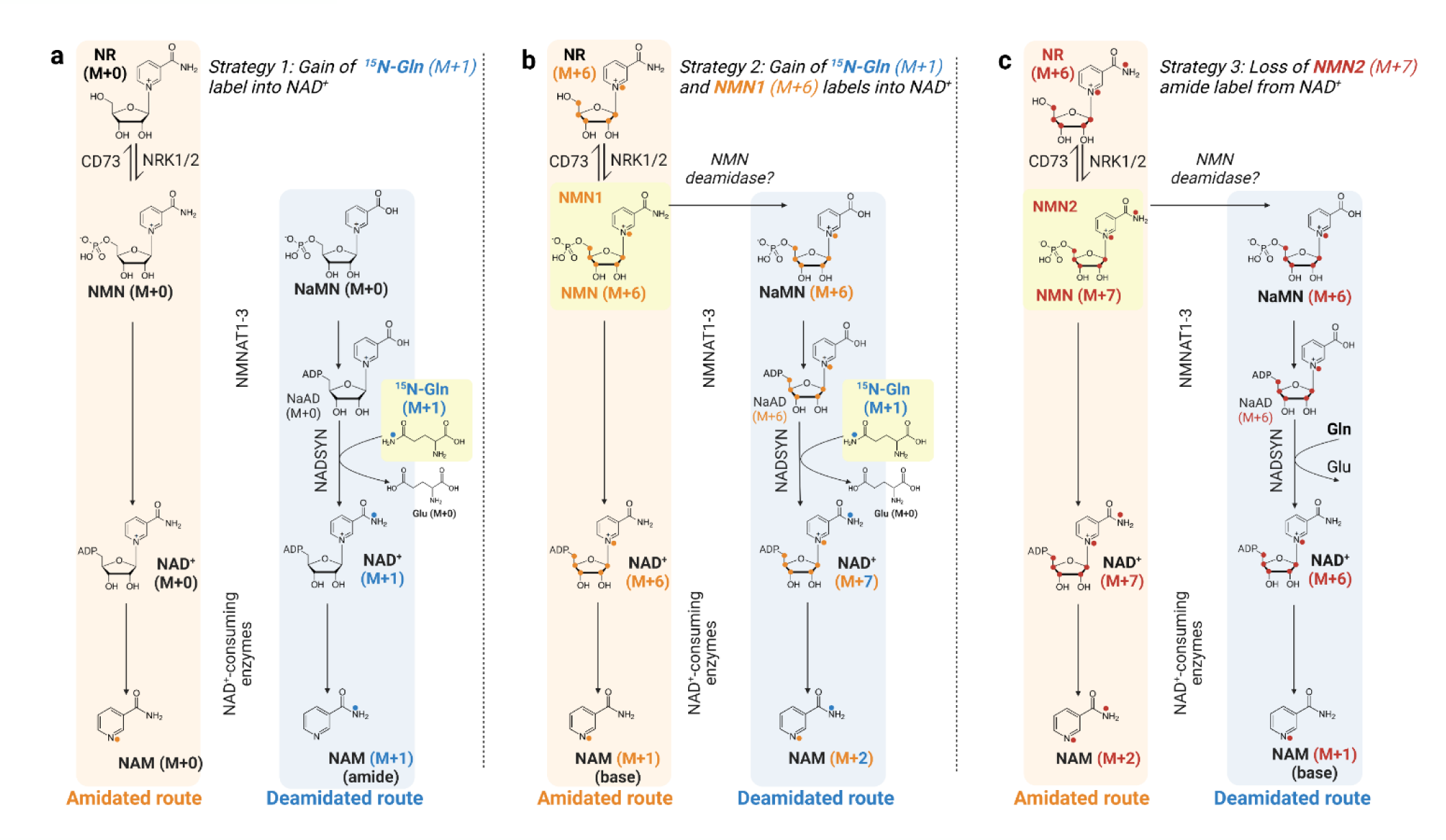
Schemes for measuring NMN incorporation in the NAD^+^ metabolome via amidated versus deamidated routes. In strategy 1 (a) ^15^N-glutamine (M+1) is provided as a substrate for the enzyme NAD synthetase (NADS), which substitutes the amide ^15^N atom from Gln for an O atom in the deamidated precursor nicotinic acid adenine dinucleotide (NaAD) to yield NAD^+^ with a ^15^N label at the nicotinyl amide position, for an overall mass shift of M+1. To assess whether NMN can be deamidated prior to its incorporation into NAD^+^, in strategy 2 (b) the isotopologue NMN1 (M+6), containing labels in the ribose and the base position of the nicotinyl ring is delivered in the presence of ^15^N-Gln as above. Incorporation of NMN1 (M+6) via the recycling route results in M+6 labelling of NAD^+^, whereas prior deamidation of NMN1 and amidation by NADS will result in an additional ^15^N label (M+1) from ^15^N-Gln, resulting in mass shift from M+6 to M+7. In scheme c) the isotopologue NMN2 is labelled at the same positions as NMN1, with an additional ^15^N label at the nicotinyl amine for an overall mass shift of M+7. Direct incorporation of NMN2 via the recycling route results in M+7 labelling of NAD^+^, whereas NMN deamination will involve loss of the additional nicotinyl ^15^N amide label, resulting in M+6 rather than M+7 labelling of NAD^+^. Following NAD^+^ biosynthesis, the expected ^15^N labelling of nicotinamide (Nam) at the amide and base positions is indicated.

While ^15^N-Gln labelling should allow for labelling of NAD that is synthesised via the enzyme NADS, this strategy does not reflect the fate of exogenous NMN. In the second strategy (Fig. 2b), we designed an isotopologue of NMN (herein designated as **NMN1**) that was ^13^C labelled at all five carbon positions of the ribose moiety for an M+5 mass shift, and ^15^N labelled at the nicotinyl ring for an overall M+6 mass shift (Fig. 2b, Supp. Fig. 2). This isotope was to be delivered in the presence of ^15^N-Gln, as above. When incorporated into NAD via the recycling pathway, **NMN1** would be expected to yield a mass shift of M+6. If **NMN1** undergoes deamidation, its incorporation into NAD^+^ would require re-amidation by NADS, resulting in the incorporation of an additional M+1 label from ^15^N-Gln, for an overall mass shift of M+7. By comparing the ratio of M+7 to M+6 NAD, this strategy would reveal the proportion of NMN that was incorporated via prior deamidation. The use of multiple reaction monitoring (MRM) mass spectrometry for targeted fragmentation and monitoring of metabolites would further reveal whether the incorporation of this extra label occurred at the nicotinyl amide position, providing further confidence in this labelling strategy.

In the third strategy (Fig. 2c), we complemented this approach using an isotope of NMN (herein designated as **NMN2**) that was labelled at the same positions as **NMN1**, with an additional ^15^N label at the nicotinyl amide group, for an overall mass shift of M+7. If **NMN2** undergoes deamidation, this additional ^15^N label at the nicotinyl amide group should be lost and substituted with an unlabelled N atom from the endogenous Gln pool, resulting in a loss of the overall mass shift from M+7 to M+6 labelling of NAD. As with the previous strategy, comparison of the ratio of M+7 to M+6 labelled NAD^+^ should reflect differences in the degree to which exogenous **NMN2** undergoes deamidation prior to its incorporation into NAD^+^.

In each of these strategies, differences in the isotopic labelling of NAD^+^ should also be reflected by comparing the labelling of nicotinamide (Nam), which is released either through NAD breakdown by NAD consuming enzymes, or could alternatively reflect the hydrolysis of exogenous NMN into free Nam by enzymes such as CD38. These Nam isotopes would be expected to be recycled back into NMN, NR and NAD^+^ (Fig. 2). An important qualifier in the interpretation of recycled intermediates is that differences in labelling may reflect the deamidation of Nam, rather than the deamidation of NMN, as has been described by others^44^. In all experiments, the use of triple-quad mass spectrometry and multiple reaction monitoring (MRM) for targeted metabolomics could further refine these data to determine where mass shifts occurred, including whether Nam was labelled at the pyridine ring (base) or amide positions, and whether M+6 or M+7 labelling of NAD^+^ was from the NMN rather than the AMP moiety.

### *In vitro* evidence for mammalian NMN deamidation in hepatocytes

To test whether this scheme would lead to labelling of the NAD pool as anticipated (Fig. 2), we first used primary rat hepatocytes grown *in vitro*, to avoid contributions from the microbiome. Hepatocytes were treated for 24 hr with ^15^N-glutamine (M+1) (Fig. 3a) in the presence or absence of **NMN1** (M+6) (Fig. 3b), or with **NMN2** (M+7) (Fig. 3c). Cell lysates were subject to targeted metabolomic analysis to assess the degree of isotope incorporation into each metabolite (Fig. 3d-k, Supp. Fig. 3). Delivery of each of these isotopes yielded the expected M+6 and M+7 mass shifts of NMN (Fig. 3e, Supp. Fig. 3b) as well as its de-phosphorylated counterpart NR (Fig. 3d, Supp. Fig. 3a), which is consistent with the indirect transport of NMN^27^. Given these NMN isotopes did not include a label on the phosphate group, these data do not however exclude the direct transport of NMN via the putative transporter SLC12A8^47^. For this reason, the data in this investigation could be interpreted as evidence for deamidation of NMN and/or NR, rather than NMN alone. As expected, delivery of ^15^N glutamine led to the expected labelling of Gln (Fig. 3f, Supp. Fig. 3e).

**Figure 3.**
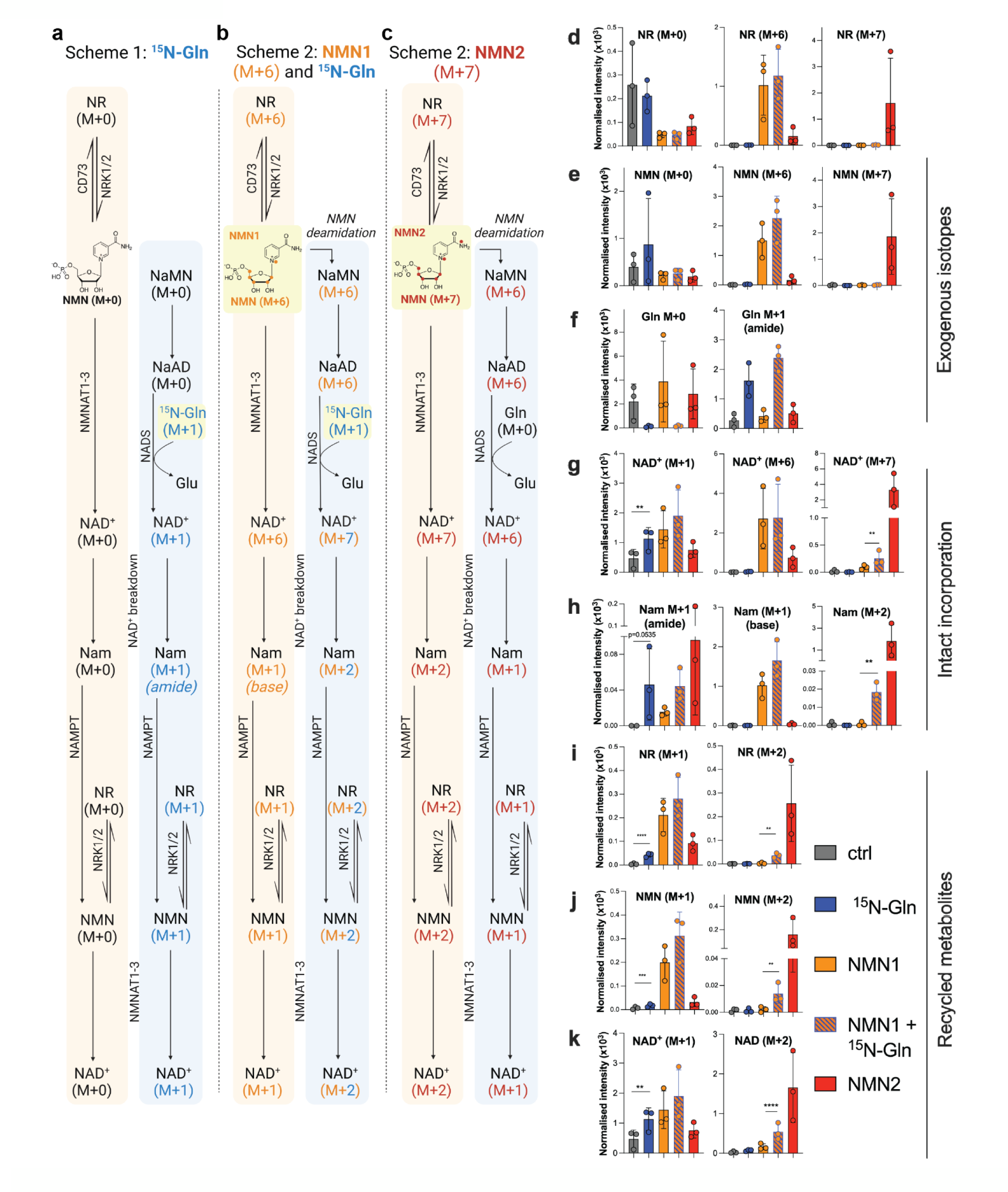
^15^N-glutamine labelling of NAD^+^ synthesis. Primary hepatocytes were treated with a) amide labelled ^15^N-glutamine (M+1) or unlabelled Gln (4 mM) alone or b) in the presence of NMN1 (M+6), or with c) NMN2 (M+7) isotopes (200 µM) alone for 24 hr, to measure the incorporation of exogenous NMN via the *de novo* pathway as described in Figure 2. ^15^N-Gln (M+1), **NMN1** (M+6) and **NMN2** (M+7) isotopes led to the expected isotopic labelling of d) NR, e) NMN and f) Gln, with subsequent incorporation into g) NAD^+^, where the formation of M+7 labelled NAD^+^ during treatment with **NMN1** (M+6) and ^15^N-Gln (M+1) is indicative of NMN incorporation via NADS. NAD^+^ consumption leads to the formation of h) free nicotinamide (Nam), with expected M+1 amide labelling by ^15^N-Gln, M+1 base labelling by NMN1 and M+2 labelling by the combination of NMN1 and ^15^N-Gln indicative of deamidation. Labelled Nam is recycled back into i) NR, j) NMN and k) NAD^+^. Data analysed by Kruskal-Wallis ANOVA with post-hoc test, n=3 biological replicates.

Next, we measured the incorporation of these labels into NAD (Fig. 3g, Supp. Fig. 3c). High levels of M+1 NAD^+^ labelling (Fig. 3g) were observed in samples treated with **NMN1**, likely due to recycling of the M+1 labelled Nam moiety following the breakdown of either NAD^+^ or NMN. As expected, treatment with **NMN1** (M+6) and **NMN2** (M+7) led to M+6 and M+7 labelling of NAD^+^ (Fig. 3g, Supp. Fig. 3c). Interestingly, we observed that ^15^N-Gln co-treatment with **NMN1** (M+6) resulted in the formation of M+7 labelled NAD^+^ when compared to **NMN1** (M+6) alone (Fig. 3g), supporting earlier observations^38^ that mammalian liver may possess NMN de-amidase activity. In line with the expected recycling of labelled Nam from NAD^+^ (Fig. 2b), this increased formation of M+7 labelled NAD^+^ during **NMN1** (M+6) and ^15^N-Gln (M+1) co-treatment was matched by an identical increase in M+2 labelling of free Nam (Fig. 3h, Supp. Fig. 3d), which was re-incorporated into the nicotinyl moiety of NR (Fig. 3i), NMN (Fig. 3j) and NAD^+^ (Fig. 3k). ^15^N-Gln treatment increased M+1 labelling at the amide position of Nam (Fig. 3h), but not the base N atom of the pyridine ring (Nam_base_, Fig. 3h, Supp. Fig. 4b), which does not undergo substitution by NADS, with **NMN1** treatment (Fig. 2a) treatment serving as a positive control for labelling at this position. These labels were incorporated into NAD at the expected positions, with subsequent MS fragments showing labelling at the nicotinyl portion of NAD, and not at the AMP portion of the overall molecule (Supp. Fig. 4a). Overall, these data verified our system of labelling, validated the specificity of our targeted bioanalytical approach, and together, point to baseline endogenous NMN de-amidase in mammals.

**Figure 4.**
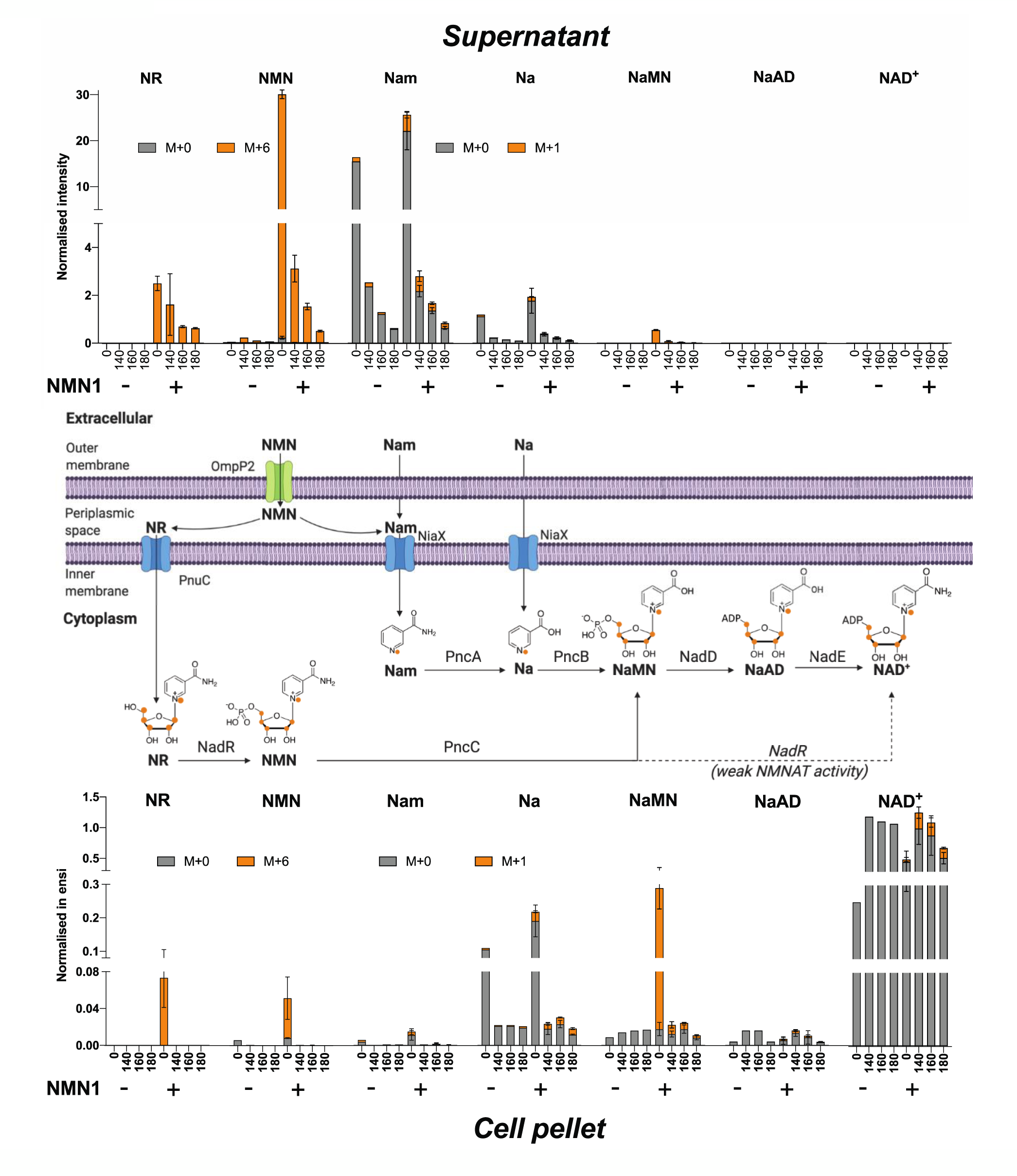
NMN deamidation in bacteria. Liquid cultures of *E. coli* OP50 bacteria were supplemented with M+6 labelled NMN1 (0.1 mM) at inoculation of a fresh culture. Samples were taken at time 0 (after NMN), 140, 160 and 180 minutes after NMN supplementation. Following separation of the culture supernatant (top) from the cell lysate (bottom), metabolites were extracted and subjected to targeted LC-MS/MS mass spectrometry to detect the incorporation of the M+6 isotope label into NMN, NR, NaMN and NAD^+^ as well as M+1 labelling of nicotinamide (Nam) and nicotinic acid (Na) in both the culture supernatant (top) and cell lysates (bottom). Data represents mean ± s.d. (n=3-5 samples per time point).

### NMN deamidation by bacteria

While experiments in isolated hepatocytes (Fig. 3) demonstrated the presence of endogenous de- amidase activity, given the magnitude of the impact of antibiotics treatment on the deamidated pathway *in vivo* (Fig. 1), we next sought to measure the incorporation of labelled NMN into the NAD^+^ metabolome of bacteria. In contrast to mammals, bacteria encode well characterised de-amidase enzymes for NMN^40^ and Nam^48^, and the gut microbiome has also been implicated in the deamidation of orally administered Nam and NR^44, 45^. This is likely due to intrinsic differences in DNA repair which is fuelled by NAD^+^ in bacteria, rather than by ATP as in mammals. This NAD^+^ dependent DNA ligase is inhibited by NMN, the product of its own reaction^41–43^, resulting in the accumulation of intracellular NMN during exponential growth^49^. This NMN is salvaged through the bacterial NMN de-amidase PncC, yielding NaMN as a substrate for NAD^+^ synthesis by the Preiss-Handler pathway^40^. To model whether extracellular NMN would undergo deamidation by bacteria, growth phase *E. coli* cultures were supplemented with **NMN1** (M+6) (Fig. 2b) and subjected to targeted metabolomics of both cell lysates and extracellular culture media (Fig. 4). Consistent with the role of PncC in NMN metabolism in bacteria, treatment with labelled NMN resulted in the rapid incorporation of isotope labels into NaMN, with vastly increased labelling of NaMN compared to NMN (Fig. 4). Similarly, a role for the Nam de-amidase PncA^44^ is strikingly reflected in the abundance of nicotinic acid (Na) compared to nicotinamide (Nam) in the cell pellet compared to the culture supernatant, where the ratio of Nam to Na in growth media was completely reversed. Overall, the avid uptake of NMN, followed by its rapid shunting into deamidated metabolites such as NaMN (Fig. 4) supports the idea that the gut microbiome could contribute to the metabolism of orally administered NAD precursors such as NMN.

### Antibiotic treatment alters NMN de-amidation *in vivo*

To directly trace whether the increase in deamidated metabolites following NMN administration (Fig. 1) was indeed due to the direct deamidation and incorporation of these metabolites, we next delivered our strategically designed isotopes into animals that had similarly been treated with antibiotics to deplete the gut microbiome (Fig. 1b, Supp. Fig. 1). Following antibiotic treatment, animals received a single oral gavage (50 mg/kg) of the **NMN1** (M+6) isotope in parallel with an i.p. bolus of ^15^N-Gln (M+1), as illustrated in Fig. 2b. As a control for labelling, animals also received ^15^N-Gln (M+1) alone, as illustrated in Fig. 2a. Four hours later, animals were sacrificed and tissues rapidly preserved for targeted metabolomic analysis (Fig. 5a-d, Supp. Figs. 5-7). In a separate experiment, a different cohort of antibiotic treated animals (Supp. Fig. 1) received a bolus of the **NMN2** (M+7) isotope (Fig. 2c) alone, following which tissues were similarly collected 4 hr later for targeted metabolomic analysis (Fig. 5e-h, Supp. Figs. 7, 8). Total levels of unlabelled and labelled metabolites are summarised in Fig. 6, with raw data in Supp. Figs. 5-8.

**Figure 5.**
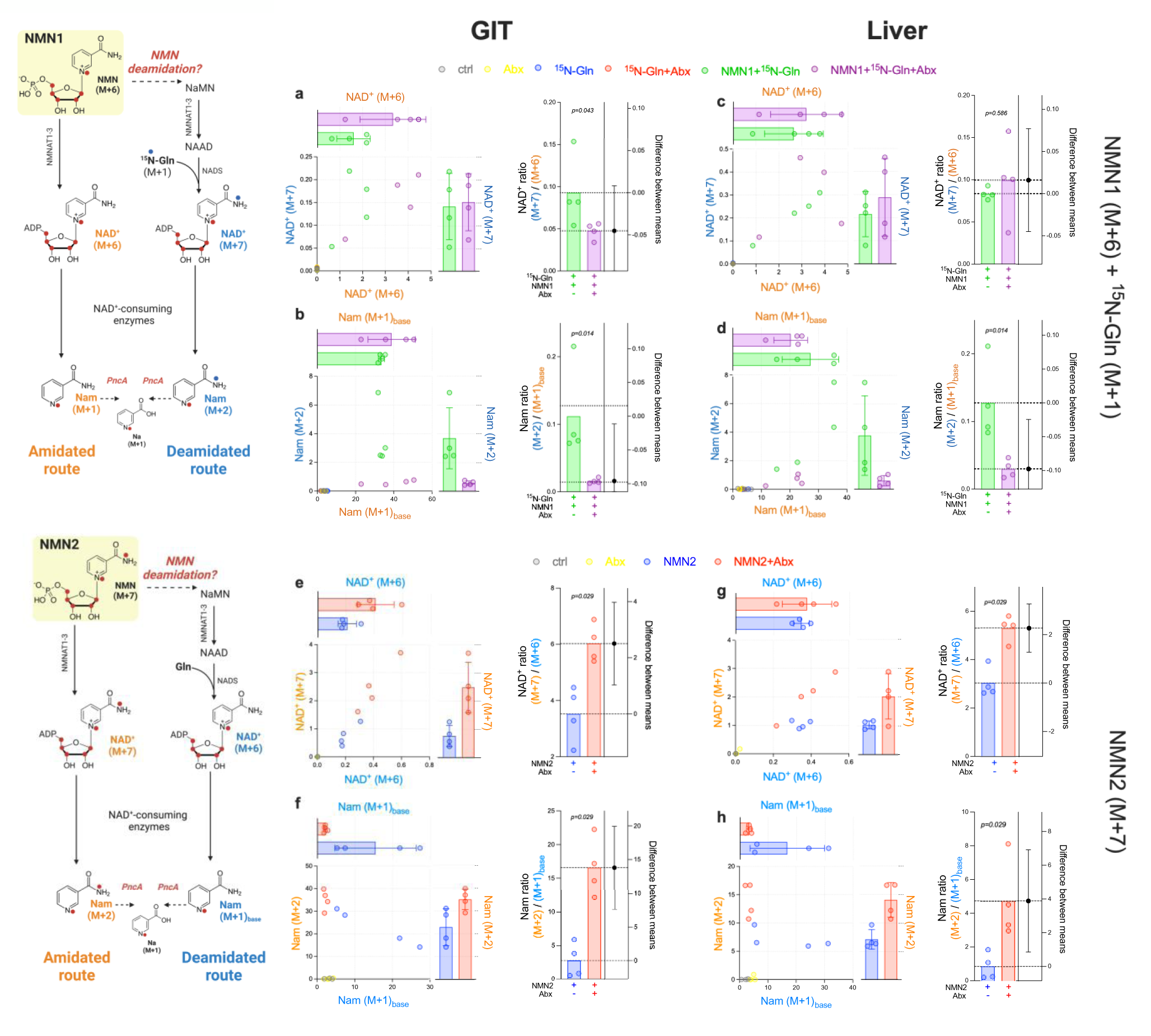
Contribution of the microbiome to NMN deamidation *in vivo*. Antibiotic (Abx) treated animals were orally administered with (a-d) **NMN1** (M+6) isotope in the presence or absence of ^15^N- glutamine (M+1), with the formation of M+7 labelled NAD^+^ or M+2 labelled Nam reflective of incorporation following deamidation and reamidation by the enzyme NADS. Antibiotic treatment reduced (a, b) M+7 labelling of NAD^+^ and (c, d) M+2 labelling of nicotinamide (Nam), which would reflect their incorporation following deamidation. Data are expressed as ratios to M+6 NAD^+^ and M+1_base_ Nam, which are the expected isotope products of NMN or NR assimilation via the canonical (amidated) route. In a separate cohort, animals were (e-h) treated **NMN2** (M+7), where loss of the amide ^15^N label to form M+6 labelled NAD^+^ or M+1_base_ labelled Nam would reflect deamidation. Antibiotic treatment protected M+7 NAD^+^ (e, g) and M+2 Nam (f, h) against loss of the NMN2 ^15^N amide compared to untreated animals. Samples in the left column (a, b, e, f) were from the gastrointestinal tract (GIT), samples on the right (c, d, g, h) were from liver. **NMN1** (a-d) and **NMN2** (e-h) experiments were run in separate cohorts of animals, each measurement represents tissue from a separate animal. Comparisons of isotope ratios between animals treated with or without antibiotics are shown as estimation plots for the difference between means, with error bars representing 95% confidence intervals. P-values represent the result of permutational t-tests, n=4 animals per group.

**Figure 6.**
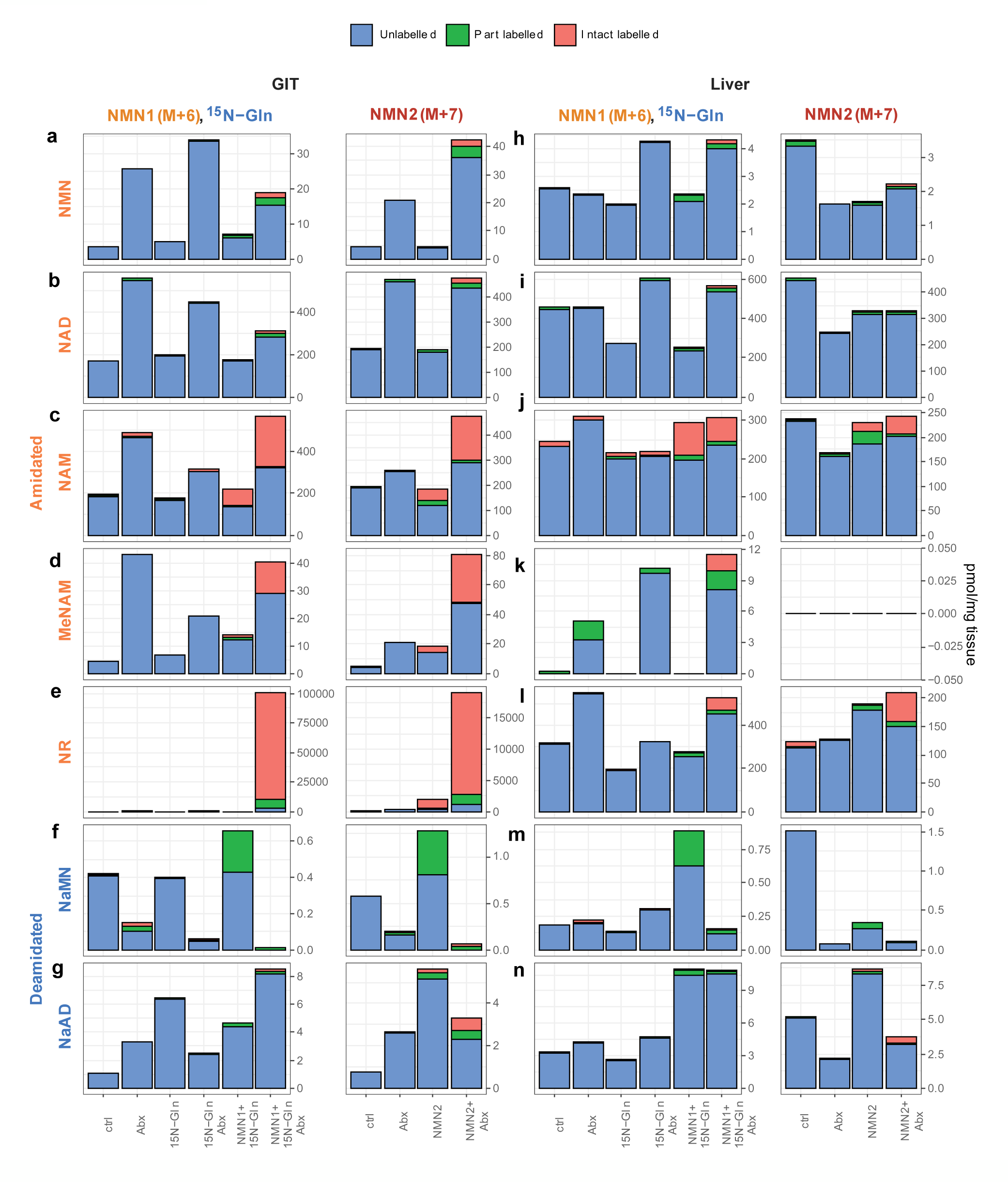
NMN1 and NMN2 label incorporation *in vivo*. Total metabolite levels in (a-g) GIT and (h-n) liver for the amidated species (a, h) NMN, (b, i) NAD^+^, (c, j) nicotinamide, (d, k) 1-methyl- nicotinamide, (e, l) NR, and the deamidated species (f, m) NaMN and (g, n) NaAD. Data are separated into each of two independent experiments using either **NMN1** (M+6) with ^15^N-glutamine or **NMN2** (M+7) alone, with colours indicating the levels of unlabelled (i.e. M+0) metabolites, intact labelled or part labelled, with the latter indicating label incorporation following the expected recycling of the initial NMN labels as shown in Fig. 3. Further details for each species are shown in supplementary data. n=4 animals per group.

**Figure 7.**
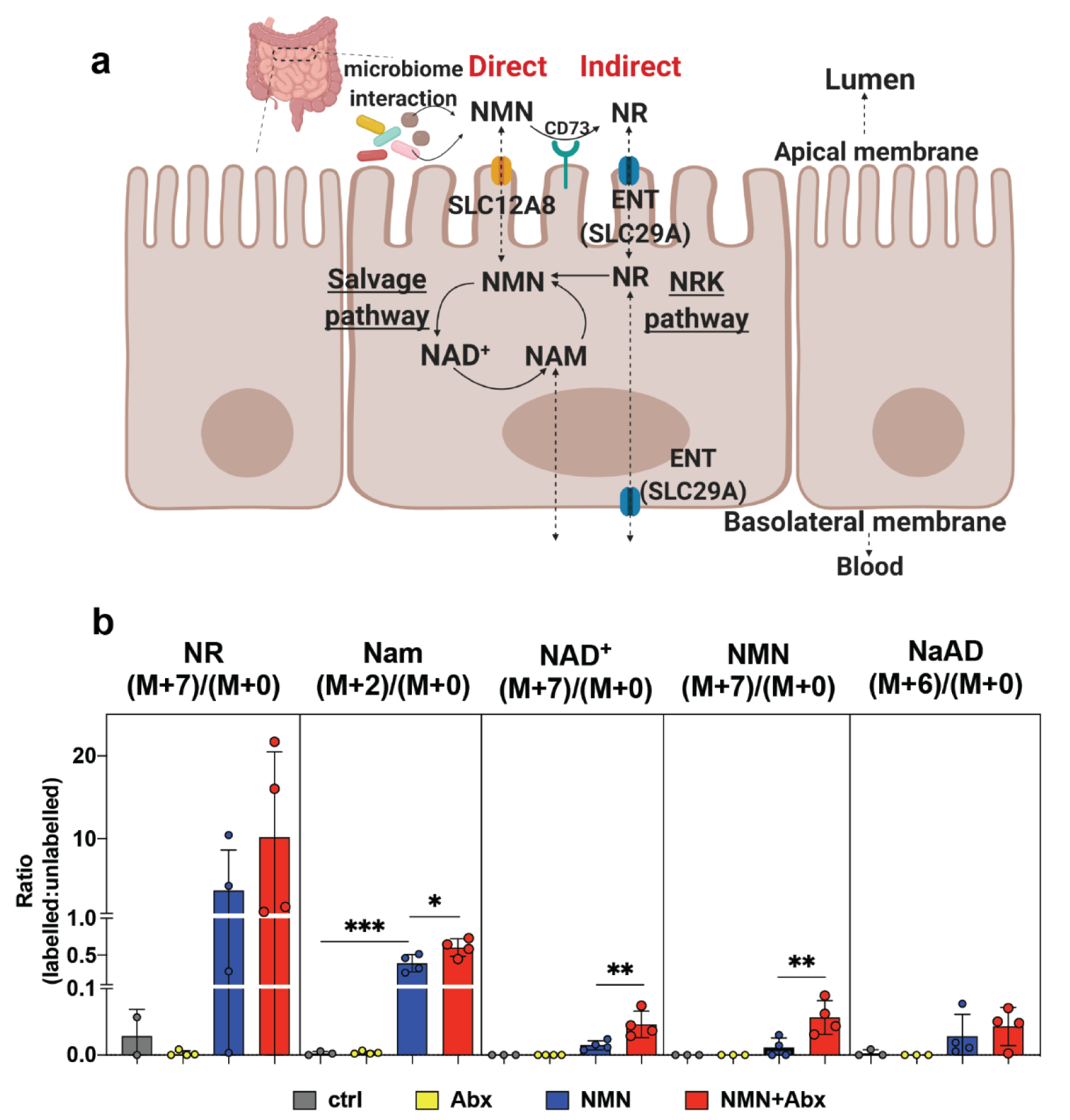
Isotope labelled NMN treatment results in greater labelling of the NR than NMN pool, suggesting indirect uptake. (a) The two proposed mechanisms for NMN uptake are either directly through the putative NMN transporter SLC12A8, or indirectly by dephosphorylation into NR via the ecto-5’-nucleotidase CD73 which is present on the apical side of intestinal cells. To compare the contributions of either direct or indirect transport mechanisms, the contribution of isotope labelled NMN to the overall pool of each metabolite is shown for the intestinal tissue of the NMN2 cohort (data from Fig. 6). Error bars are s.d., each data point represents tissue from a separate animal.

In tissues from animals treated with **NMN1** (M+6), the deamidation of NMN could be quantified by comparing the ratio of M+6 NAD^+^, which would assume incorporation following the canonical route, to M+7 NAD^+^, which had incorporated an extra mass shift from co-treatment with ^15^N-Gln (M+1) (Fig. 2b). In this experiment, an increased ratio of M+7 to M+6 labelled NAD^+^ would indicate the deamidation of NMN. The reason for using ratios, rather than the overall amounts of each isotope (Fig. 6b, Supp. Fig. 5), is that they internally control for differences in bioavailability within each animal. From the intact labelling of NAD^+^ in the GIT from **NMN1** treatment, around 13% was M+7 labelled (Fig. 5a). Importantly, these data likely underestimate incorporation via the deamidated route, as this scheme relied on the availability of exogenous ^15^N-Gln relative to the endogenous pool of unlabelled Gln, which composed only 8-12% of the total plasma Gln pool at the 4 hr timepoint (Supp. Fig. 6a). Consistent with our hypothesis, the ratio of M+7 to M+6 labelling in the GIT was reduced in antibiotic treated animals (Fig. 5a), suggesting reduced deamidation of orally administered NMN when contributions from the microbiome were reduced. This was reflected by a reduction in the ratio of M+2 to M+1_base_ labelled Nam (Fig. 5b), however this change also likely reflected reduced contributions from the bacterial nicotinamide de-amidase PncA^48^ following antibiotic treatment^44^. A similar trend was observed in both GIT (Fig. 5a, b), liver (Fig. 5c, d) and plasma (Supp. Fig. 7).

To complement this approach, a separate cohort of animals previously treated with antibiotics were treated with **NMN2** (M+7) (Fig. 5e-h, Supp. Fig. 7-8). In this strategy (Fig. 2c), we would anticipate that de-amidation by the microbiome would result in loss of the ^15^N amide label, resulting in the formation of M+6 NAD^+^ at the expense of M+7 NAD^+^. In contrast to the previous **NMN1** experiment, the ratio of M+7 to M+6 NAD^+^ would instead decrease as the rate of de-amidation increased. Further, interpretation of deamidation in this **NMN2** experiment was not limited by the availability of exogenous ^15^N-Gln relative to a large, endogenous pool of Gln, as was the case with the **NMN1** experiment (Fig. 5a-d, Supp. Fig. 6). In this experiment, the ratio of M+7 to M+6 NAD^+^ was around 3:1 (Fig. 5e), suggesting that around 25% of orally administered NMN undergoes deamidation prior to its intact incorporation into NAD^+^. In agreement with the previous experiment, the ratio of M+7 to M+6 labelled NAD^+^ was increased in the GIT and liver of antibiotic treated animals (Fig. 5e). This was similarly matched by an increased ratio of M+2 to M+1_base_ labelled Nam (Fig. 5f) in antibiotic treated animals, reflecting decreased incorporation following deamidation, though this could instead be due the deamidation of Nam rather than NMN. Similar trends in these ratios were observed for both GIT (Fig. 5e, f) and liver (Fig. 5g, h). In addition to differences in the isotope labelling of NAD^+^ (Fig. 5), these experiments replicated the inverse relationship between NaMN and NMN levels following antibiotic treatment (Fig. 6a, f) observed in our earlier experiment with unlabelled NMN (Fig. 1n). Overall, data from these two complementary *in vivo* isotope labelling experiments support the concept that orally delivered NMN or NR can undergo de-amidation prior to incorporation, and a role for the microbiome in mediating this. While these data could in part explain the spike in the de-amidated metabolites NaMN and NaAD following treatment with the amidated precursors NR^26^ or NMN (Fig. 1), it is important to note that when measured as a proportion of the overall NAD^+^ pool, the contribution of both M+7 and M+6 intact labelled NAD^+^ was small. Partially labelled NAD^+^ (M+2) was around 10-fold more abundant than intact labelled NAD^+^ (M+7) (Fig. 6b, Supp. Figs. 5, 8), indicating either cleavage of the labile glycosidic bond of NMN prior to its incorporation, or rapid recycling of NAD^+^^50^. Following cleavage of the glycosidic bond to release free Nam, its deamidation in the GIT^51, 52^ by the bacterial enzyme PncA^44^ also likely contributes to these changes.

In both experiments, the only NAD^+^ metabolite reliably detected in plasma samples was nicotinamide (Supp. Fig. 6), where isotope labelling for each experiment was in line with the results for other tissues.

### Host-microbe interactions in the bioavailability of orally delivered NAD^+^ precursors

Another unexpected aspect of these data was the overall increase in levels of these metabolites due to antibiotics treatment alone, which more than doubled the labelling of the metabolites NMN, NR, NAD^+^ and Nam (Fig. 1i-k, 6a-e). This increase even occurred in unlabelled metabolites in animals that did not receive exogenous NMN (Fig. 6a-d). When either **NMN1** (M+6) or **NMN2** (M+7) were delivered, the incorporation of exogenous labels into NAD^+^ metabolites was vastly increased in antibiotics treated animals, in the case of NR in the gut, by an order of magnitude (Fig. 6e). A similar trend was observed for the increase in NR levels in antibiotics treated animals that received unlabelled NMN (Fig. 1i). The overwhelming abundance of NR as the dominant NAD metabolite in the GIT, especially following NMN delivery in antibiotics treated animals is worthy of later investigation. The abundance of this single metabolite during NMN treatment in antibiotics treated animals was greater than all other NAD^+^ metabolites combined, including NAD^+^ itself. Together, the striking increase in the uptake and overall abundance of both labelled and unlabelled NAD metabolites in antibiotics treated animals suggests that the microbiome could be in competition with mammalian tissue for the uptake of orally administered, exogenous NAD precursors, and the uptake of NAD precursors from dietary sources. Future studies should measure isotope labelling of NAD^+^ metabolites in faecal contents of mice to confirm whether these compounds are being utilised by the microbiome, rather than being excreted via other mechanisms, and should use animals in which the microbiome has been reconstituted to control for the effects of antibiotics treatment.

### Evidence for NMN uptake following dephosphorylation into NR

NMN uptake can occur following the dephosphorylation of NMN into NR by the cell surface enzyme CD73, prior to uptake by ENT nucleoside transporters and re-phosphorylation into NMN inside the cell by NRK1/2 (Fig. 7)^27, 28^. Alternatively, the solute carrier protein SLC12A8 has been described as a dedicated NMN transporter^47^. As with CD73 and ENT, SLC12A8 is located on the apical side of the intestinal tissue. As both mechanisms could co-exist, the question is the degree to which each mechanism contributes to the uptake of NMN^53^. If the direct route via SLC12A8 prevailed, we would expect to see high levels of labelled (M+7 or M+6) NMN, with lesser uptake of labelled NR. In contrast, if the indirect transport of NMN following its dephosphorylation into NR was dominant, there would be a higher levels of NR labelling. In primary hepatocytes (Fig. 3), NMN1 (M+6) treatment resulted in near complete labelling of the NR pool (Supp. Fig. 3a), with slightly lesser labelling of the NMN pool (Fig. 3b). Strikingly, *in vivo* treatment showed strong labelling of the NR, but not NMN pools (Fig. 6b, e; Supp. Fig. 5b-c, Supp. Fig. 8b-c). Intact (M+6 or M+7) labelled NR levels were five-fold higher than endogenous NR (M+0), which increased to a ten-fold greater enrichment with antibiotics treatment (Fig. 7). In stark contrast, only around 5% of the NMN pool was M+7 labelled (Fig. 7). The inability of exogenous NMN to displace the endogenous NMN pool, combined with the surge of labelled NR, suggests that NMN uptake bypasses direct transport, and would instead support the dephosphorylation of NMN into NR to facilitate its intestinal absorption (Fig. 7). If direct transport of NMN does occur, its metabolism is mediated by the microbiome, as even when M+6 or M+7 labelling of NMN was observed at low levels, this only occurred in antibiotic treated animals (Fig. 6b, Supp. Fig. 6h, 8h). An important caveat of this interpretation is that limited availability of isotope labelled material meant that this study used a single time point, rather than a time course which also encompassed very early timepoints, possibly missing the minute-order kinetics of direct NMN transport that were previously reported^21, 54^.

## DISCUSSION

Together, this work provides evidence for the partial incorporation of exogenous NMN into the NAD metabolome via the deamidated route, and for contributions of the gut microbiome to the metabolism of exogenous NMN. This is in line with recent findings around the role of Nam deamidation by bacteria^44^ and the cycling of NAD precursors between the circulation and the gut microbiome^45^, however a role for the microbiome in the uptake of exogenous NMN has not been described. It will be interesting to determine whether this relationship persists for other precursors or is unique to NMN. Rather than being evidence for a “competition” relationship, differences in the uptake of exogenous NMN could reflect its role as an inhibitor of bacterial DNA ligase^41–43^, and bacterial mechanisms to prevent its accumulation. In addition to NMN de-amidase enzymes, this could include a role for bacterial NAD glycohydrolase enzymes, and/or bacterial SARM-like enzymes^55^.

While intravenous delivery of NR or NMN results in a small degree of intact assimilation into peripheral tissues such as the liver, kidney and muscle, oral delivery of NMN results in hepatic cleavage at the glycosidic bond yielding free nicotinamide due to the action of the liver^50^. Our results were in close alignment with those findings^50^, where the ratio of intact M+7 or M+6 to M+0 unlabelled NAD^+^ was around 2%, whereas the ratio of M+2 labelled to M+0 unlabelled NAD^+^, presumably as a result of incorporation of free Nam, was over 10% (Fig. 6b, Supp. Figs. 6l, 8l). Given this evidence for the decomposition of NMN into free Nam prior to its uptake, a key question for the field is why downstream precursors in NAD^+^ synthesis such as NMN and NR lead to different outcomes compared to Nam alone^56–58^. Another key question is around the cross-over between the amidated and de-amidated pathways of NAD^+^ synthesis. While we observed that exogenous NMN increased liver NaAD levels (Fig. 1e, 6g), as was previously reported for exogenous NR treatment^26^, the majority of this increase was from unlabelled NaAD (Fig. 6g). These results were similar to those of Trammel et al^26^, where isotope tracing of double-labelled NR showed that while total NaAD levels increased by over 40-fold following NR treatment, only around 45% of this NaAD was isotope labelled – with the endogenous origin of the remaining 55% remaining unexplained. These findings run against the assumed model that exogenous NAD^+^ precursors raise NAD^+^ levels through their direct incorporation into the NAD metabolome, and we suggest instead that treatment with exogenous precursors such as NMN could indirectly trigger endogenous NAD^+^ biosynthesis. The mechanism for this is not yet clear, though given the profound effect of antibiotic treatment, especially the overwhelming abundance of NR in the gut (Fig. 6e, Supp. Fig. 6b, 8b), could involve interplay with the gut microbiome. One possibility for the changes in endogenous NAD^+^ metabolites following exogenous NMN treatment could be the activation of unknown signalling pathways that indirectly alter endogenous NAD metabolism, rather than the direct incorporation of exogenous material.

Another explanation is that increased substrate levels alter the *in vivo* kinetics of NAD biosynthetic enzymes, increasing the utilisation of endogenous substrates. An important question regarding NR is how its endogenous production is increased by exogenous NMN. NR is available from dietary sources^28^, and is an intermediate in the uptake of extracellular NMN^59, 60^. NMN accumulation in neurons can trigger cell death through the NADase SARM1^61^, and disposal of NMN through its adenylation into NAD^+^ can protect against neuronal death^62^. It is possible that exogenous NMN triggers pathways that degrade endogenous NMN into NR, which could act as a reservoir for NAD precursors, however the NMN ectonucleotidase CD73 that carries this out sits on the extracellular face of the plasma membrane^59, 60^ rather than the cytosol. In addition, the sheer molar quantity of unlabelled NR that we observed in the gut (Fig. 1i, 6e) relative to other metabolites challenges this idea. NAD homeostasis is tightly maintained within a defined range^63^, and the activity of NMNAT enzymes that carry out the last step of NAD biosynthesis is reversible^64^. It is possible that exogenous NAD precursors push the equilibrium of this step in the opposite direction, increasing endogenous NMN production from NAD^+^, though how intracellular NMN could be dephosphorylated to an NR reservoir in mammals is unknown. This concept of increased NAD breakdown during treatment with exogenous NMN in young animals is also supported by the increased formation of unlabelled Nam in plasma (Supp. Fig. 6a). Further, while labelled NMN treatment *in vitro* (Fig. 3) results in the formation of intact labelled NAD^+^, this occurs at the cost of unlabelled NAD^+^, for a net zero change in total NAD levels (Supp. Fig. 3c). This could suggest that when NAD^+^ is replete, the utilisation of exogenous NMN into newly synthesised NAD^+^ results in a commensurate breakdown of existing material – potentially explaining the formation of unlabelled metabolites such as NR during treatment with exogenous, labelled NMN.

Rather than degrading or recycling existing metabolites into NR, another possibility is that exogenous NMN or NR could trigger a currently unknown step in mammals that leads to endogenous NR production. While NR can be produced by the reversible phosphorolysis of Nam and ribose-5- phosphate by a purine nucleophosphorylase (PNP) in *E. coli*^65^, this step is irreversible in mammals^60, 66, 67^, and other potential steps involved in endogenous NR production in mammals are unknown. Further work is needed to understand how this occurs, for example, whether it is due to the acute up-regulation of NAD^+^ biosynthetic enzymes, the timescale by which this increase occurs, and a direct comparison of different isotope labelled NAD^+^ precursors to identify which metabolites trigger the production of endogenous metabolites under normal circumstances and during depletion of the microbiome. Regardless, the ability to trigger the production of endogenous NAD metabolites could explain why exogenous NR and NMN treatment lead to differences in pharmacokinetics, metabolite production and therapeutic effects when compared to Nam alone^56–58^, despite their rapid metabolism into free Nam by the liver^50^ (Fig. 6c, Supp. Figs. 5r, 8r).

Another speculative idea is the existence of a signalling pathway in the GIT that is sensitive to both exogenous NAD precursors and to microbial metabolites, which can mediate endogenous NAD metabolism. Metabolite sensing members of the G-protein coupled receptor (GPCR) family are putative candidates for this role as there are GPCRs already known to respond to extracellular nicotinic acid and to NAD^+^ itself^68^. One possible candidate is GPR109a, which acts as a receptor both for nicotinic acid^69^ and for butyrate, released from the microbial fermentation of dietary fibre^70^. This could link the observations including the deamidation of orally derived NAD precursors into their acid equivalents, a role for microbiome depletion in triggering the production of endogenous NAD metabolites, and evidence for the poor incorporation of intact NR or NMN into the NAD metabolome.

These conclusions are limited by using only a single timepoint for experiments, which was related to our limited availability of NMN isotopes that were expensive and difficult to produce. Recent work has identified a complex interplay between the gut microbiome and mammalian circulation in NAD homeostasis, with differing roles for each section of the gut^45^, however this study analysed the entire GIT as a single homogenised sample, potentially missing subtleties that may occur in different regions of the gut. While we observed an increase in NAD^+^ metabolite levels with antibiotics treatment, we cannot conclude that this is due to competition with the gut microbiome as we did not collect gut contents for metabolite analysis. Another key limitation is the use of antibiotics alone to ablate the gut microbiome, and further confirmation of our findings should involve experiments where the gut microbiome is reconstituted. Finally, while the isotope labelling patterns observed were largely as expected, our detection method included a small amount of isotope labelling in control samples that were not treated with exogenous isotopes, even after subjecting data to isotope correction calculations, suggesting a margin of error for these measurements.

Together, findings from these orthogonal approaches support the de-amidation of NMN by endogenous (Fig. 3) and microbial (Fig. 5) pathways prior to its incorporation into the NAD metabolome. Our findings are consistent with early descriptions of mammalian de-amidase activity^37–, 39^ and more recent work that integrates microbial and mammalian host NAD metabolism^44, 45^, though note that the direct NMN de-amidation by the gut microbiome as described here likely accounts for a small fraction of de-amidation of the overall NAD^+^ metabolome. We also conclude that NMN cellular uptake most likely occurs following its dephosphorylation into NR^27^. Regardless of this, we also note the relatively low levels of intact incorporation *in vivo* which is in line with the findings of others^50^, raising questions around the utility of NMN and NR compared to nicotinamide alone. A surprising observation of this study was the increase in the levels of both NAD^+^ metabolites in the gut with antibiotics treatment, and the increase in both labelled and unlabelled NR levels in the gut during antibiotics treatment with NMN. Future work will aim to investigate this further.

## Methods

Methods are available in Supplementary Material, with raw data available on our Mendeley data site

## Acknowledgements

Funding was from the National Health and Medical Research (NHMRC) of Australia as a Career Development Fellowship APP1122484 to LEW, and sponsored research from Jumpstart Fertility. We wish to thank anonymous donors for philanthropic support.

## Author contributions

LJK conducted experiments, analysed data, prepared figures, wrote manuscript. TJC and RM conducted microbiome analyses. EWKP, TTC, JW prepared isotope labelled NMN. SPT and DAS provided critical feedback and interpretation. LEQ conducted experiments, extracted and analysed data, wrote manuscript. LEW conceived of and designed study, obtained funding, supervised experiments, analysed data, prepared figures, wrote manuscript.

## SUPPLEMENTARY FIGURES

**Supplementary Figure 1.**
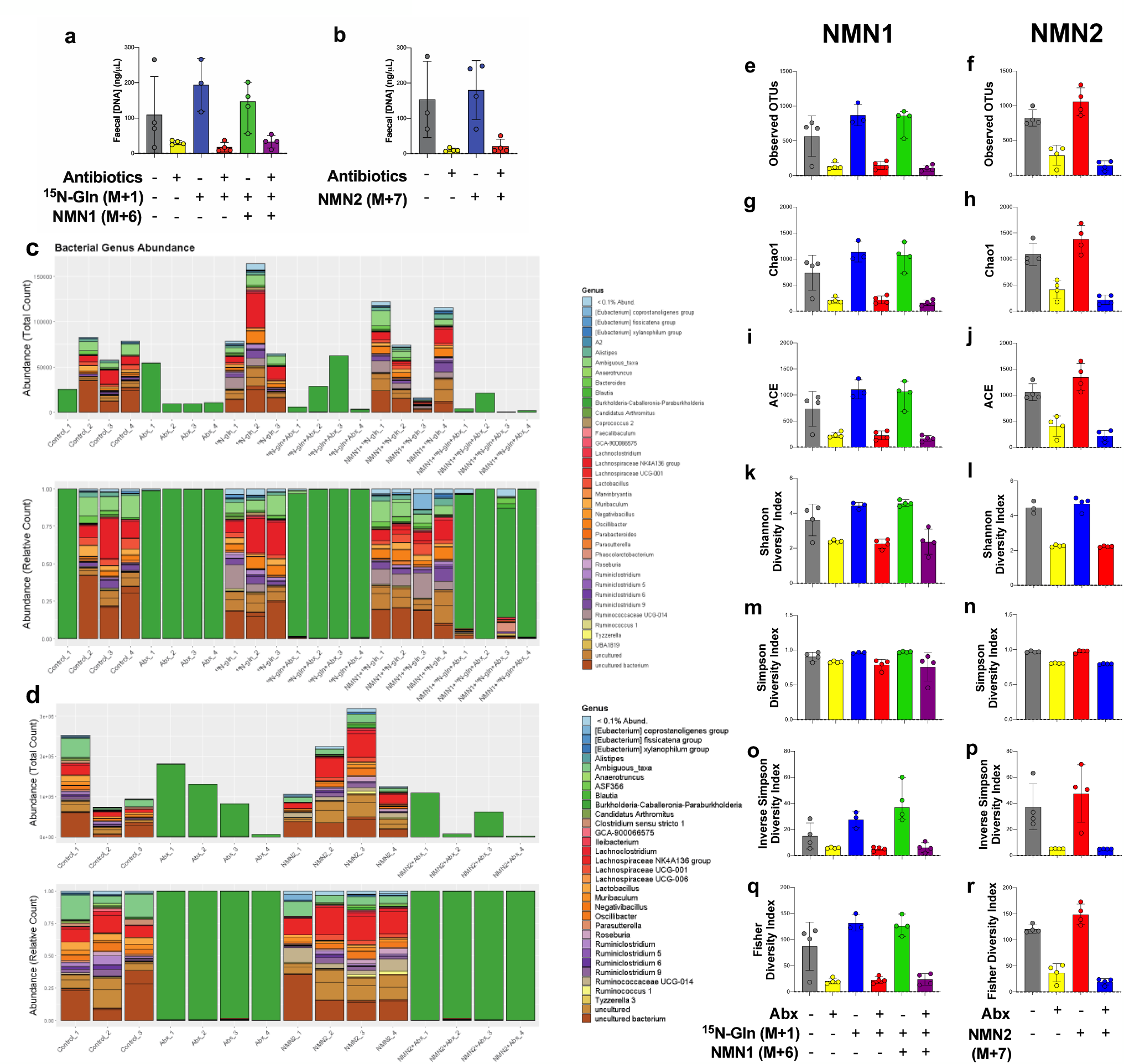
Antibiotics treatment ablates the gut microbiome. Following antibiotic treatment in the NMN1 and NMN2 mouse cohorts, (a-b) DNA was extracted from faeces to measure changes in DNA concentration. Uniform amounts of DNA were then subject to (c-d) full-length 16S rRNA Nanopore sequencing, with species abundance shown here at the genus level. Sequencing revealed a reduction in (e-f) operational taxonomic units (OTUs), the (g-h) Chao1 and (i-j) ACE species richness indices, and the (k-l) Shannon, (m-n) Simpson, (o-p) Inverse Simpson and (q-r) Fisher diversity indices. Data shown are non-rarefied; rarefication showed identical results (data not shown). Each data point represents samples from a separate animal.

**Supplementary Figure 2.**
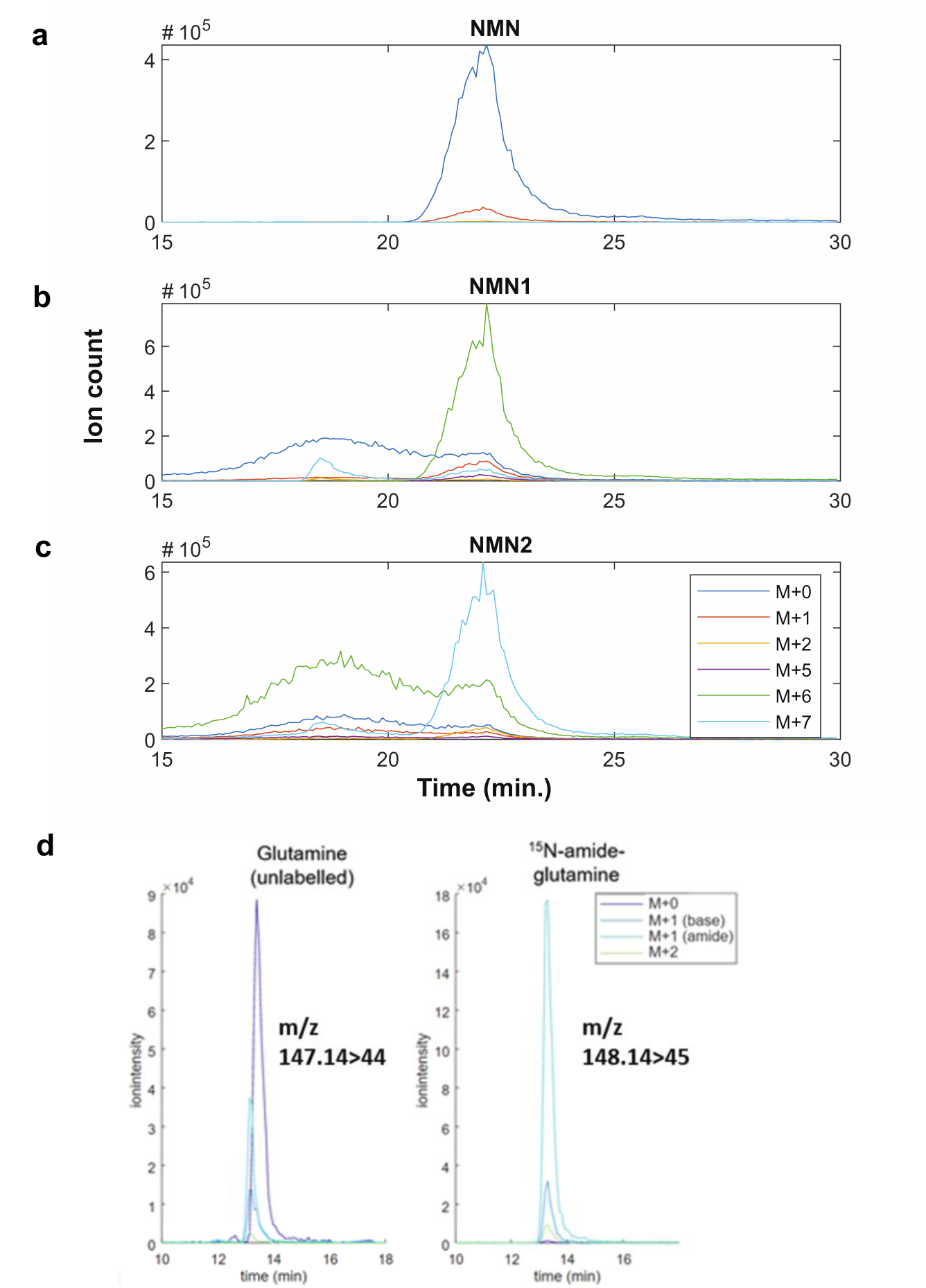
Chromatogram of isotope labelled NMN and glutamine (Gln) using MRM LC-MS/MS. The above chromatograms represent individual peaks (ion count) for a) 100µM unlabelled NMN in combined NAD metabolite standard curve mixture, b) M+6 labelled NMN1 and c) M+7 labelled NMN2, as well as d) unlabelled and ^15^N-amide labelled glutamine.

**Supplementary Figure 3.**
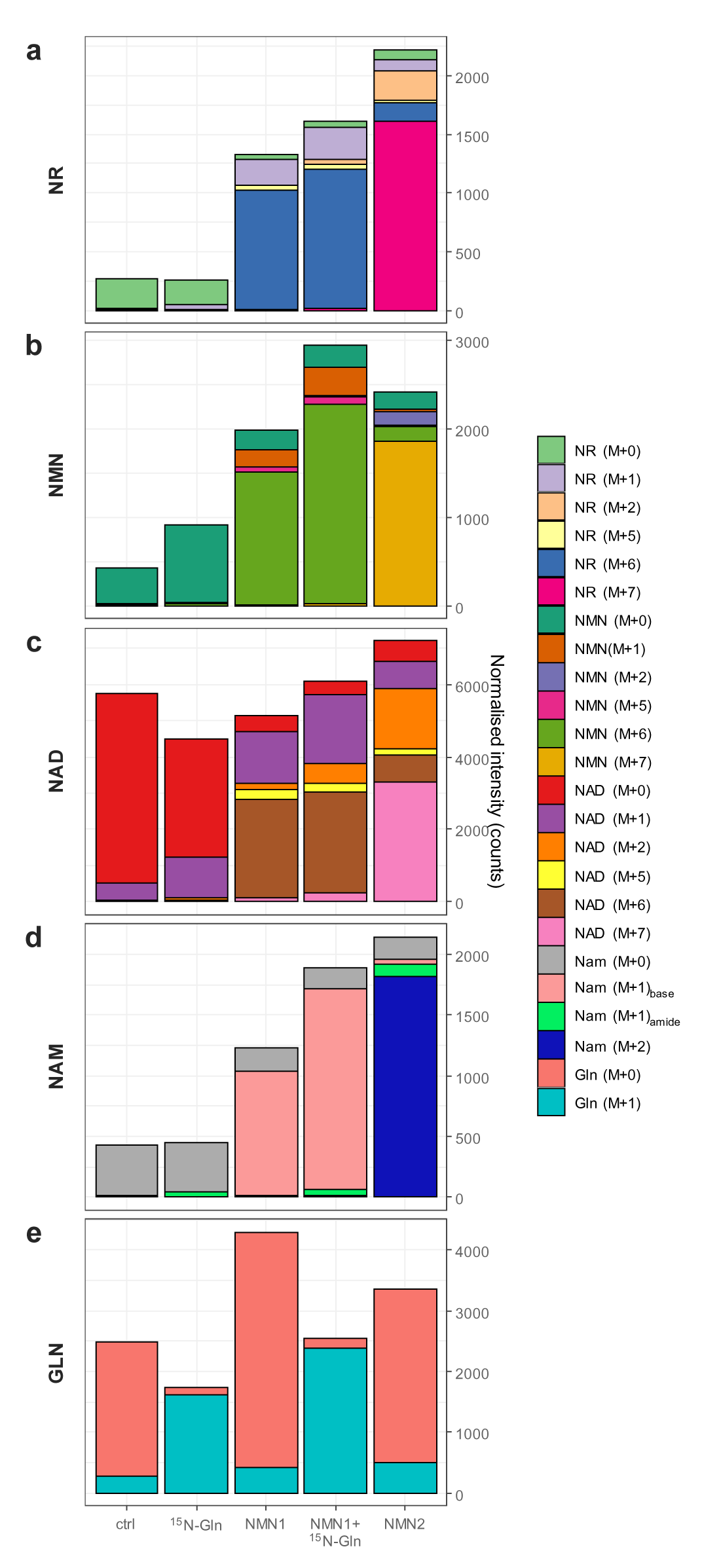
Total levels of NAD^+^ metabolites in NMN1 and NMN2 treated hepatocytes, related to main Fig. 3. Primary hepatocytes treated with ^15^N-Gln, NMN1 (M+6), or NMN2 (M+7) as in main Fig. 3. Total levels of (a) NR, (b) NMN, (c) NAD^+^, (d) nicotinamide and (e) glutamine, with contributions from each of isotope-labelled species shown.

**Supplementary Figure 4.**
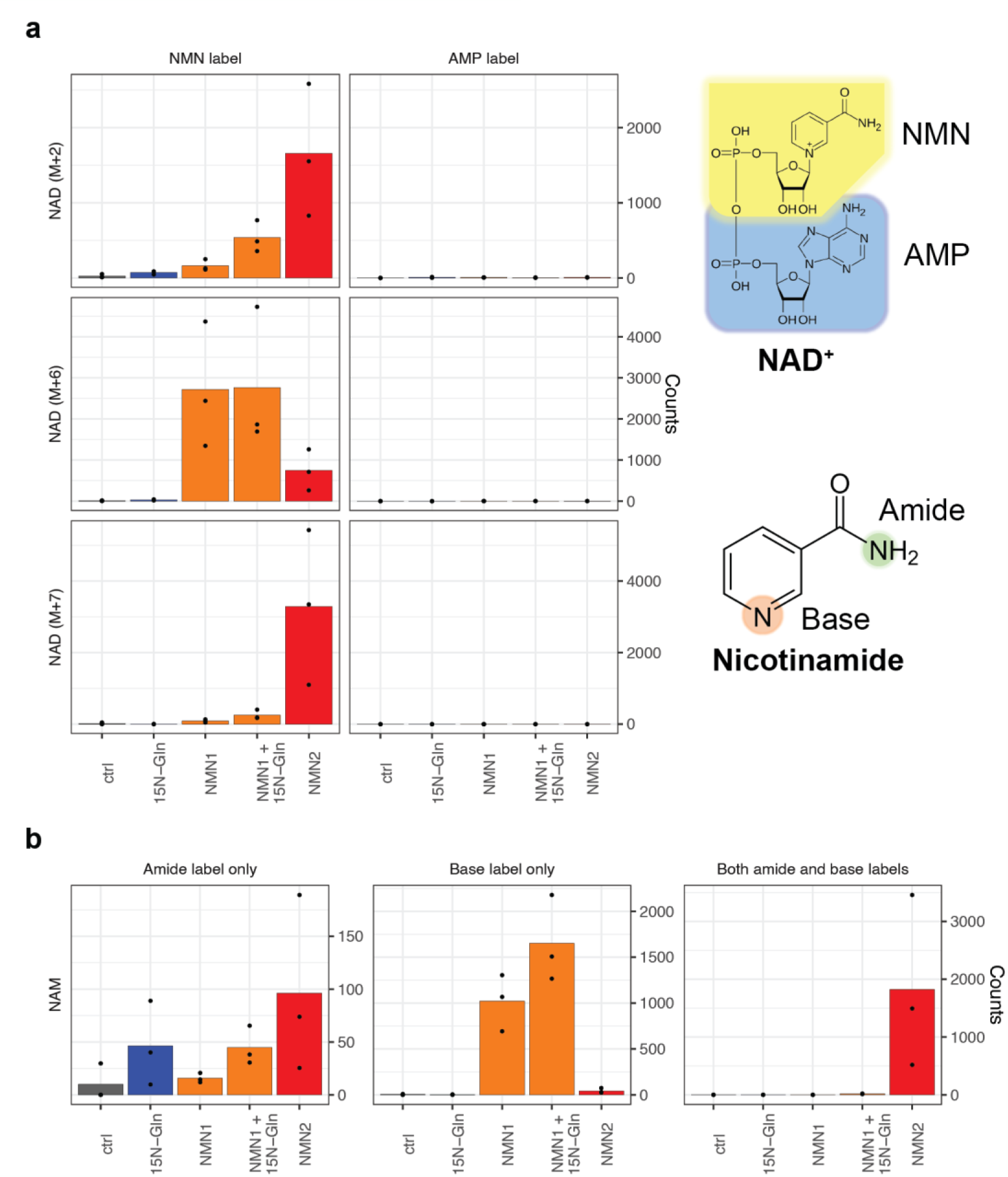
Multiple reaction monitoring (MRM) to identify fragments containing isotope labels – related to main Fig. 3. MRM enables measurement of label incorporation at (a) either the nicotinyl or adenyl fragments of NAD^+^ for each of the overall parent ion mass shifts of M+2 for nicotinamide labelling, M+6 or M+7 for both ribose and nicotinamide labelling, following treatment with NMN1 or NMN2 in primary hepatocytes as indicated. M+6 or M+7 labelling as shown in this manuscript therefore refers to label incorporation at the NMN moiety of NAD^+^, with negligible incorporation at the AMP moiety. (b) Labelling at the base (ring) or amide positions of nicotinamide for the same experiment.

**Supplementary Figure 5.**
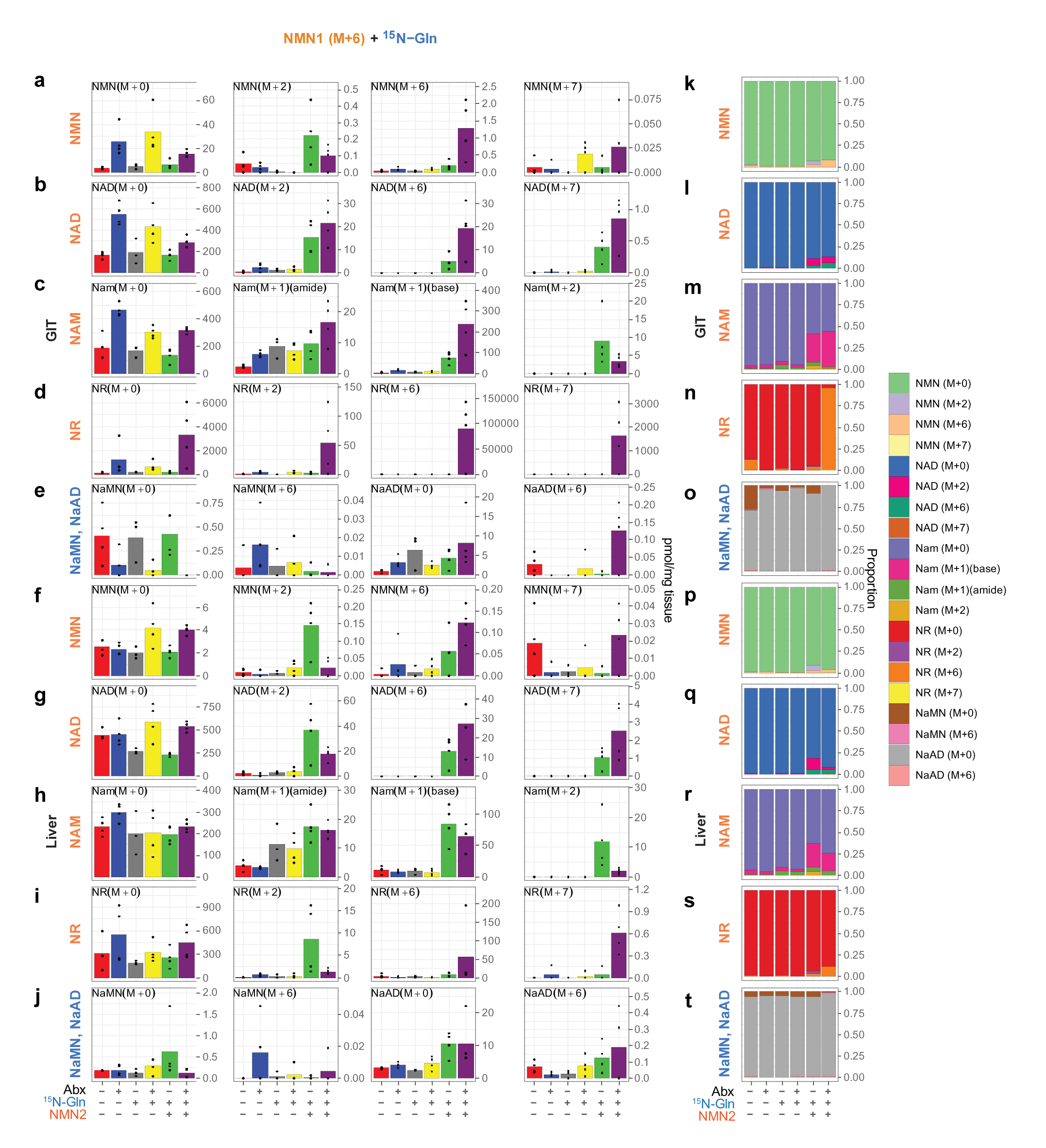
Incorporation of NMN1 (M+6) and ^15^N-glutamine (M+1) into the NAD metabolome *in vivo*. Animals were treated with antibiotics (Abx) to deplete the microbiome, followed by an oral gavage (50 mg/kg) of **NMN1** (M+6) with adjacent i.p. administration of ^15^N-Gln (M+1) (735 mg/kg, 10 ml/kg body weight). Four hours later, GIT (a-e) and liver (f-j) were collected and rapidly preserved for targeted metabolomics analysis to identify labelling of (a, f) NMN, (b, g) NAD^+^, (c, h) nicotinamide (Nam), (d, i) NR, (e, j) NaMN and NaAD, with each isotopologue shown as picomoles per mg tissue. (k-t) Data are also presented as isotope proportions of each species. n=3- 4 animals per group, raw data points shown.

**Supplementary Figure 6.**
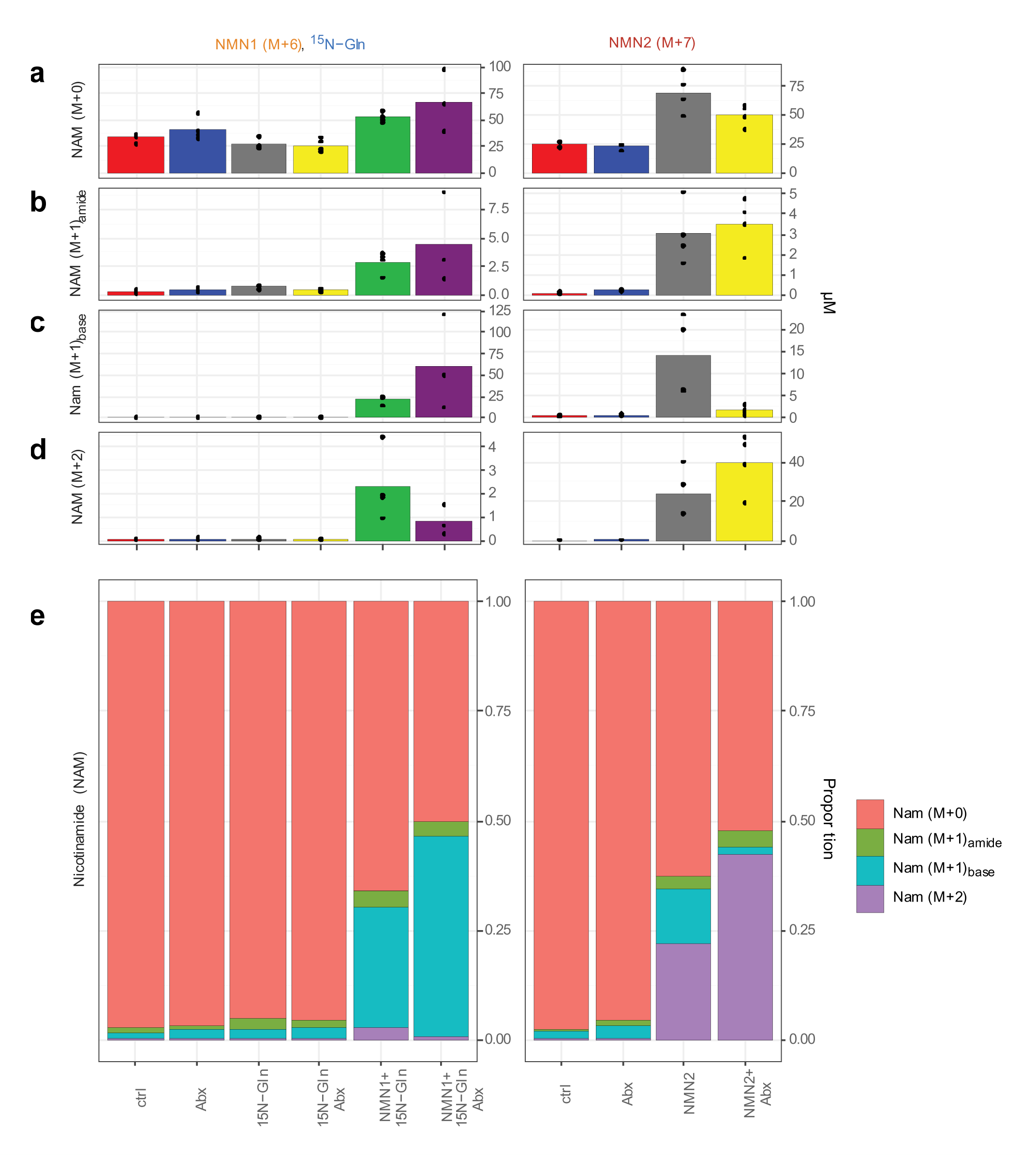
Nicotinamide (Nam) labelling in plasma. The only NAD^+^ metabolite reliably detected in mouse plasma was nicotinamide, with isotope labelling following treatment with NMN1 (M+6) and ^15^N-glutamine (M+1) or NMN2 (M+7) as indicated.

**Supplementary Figure 7.**
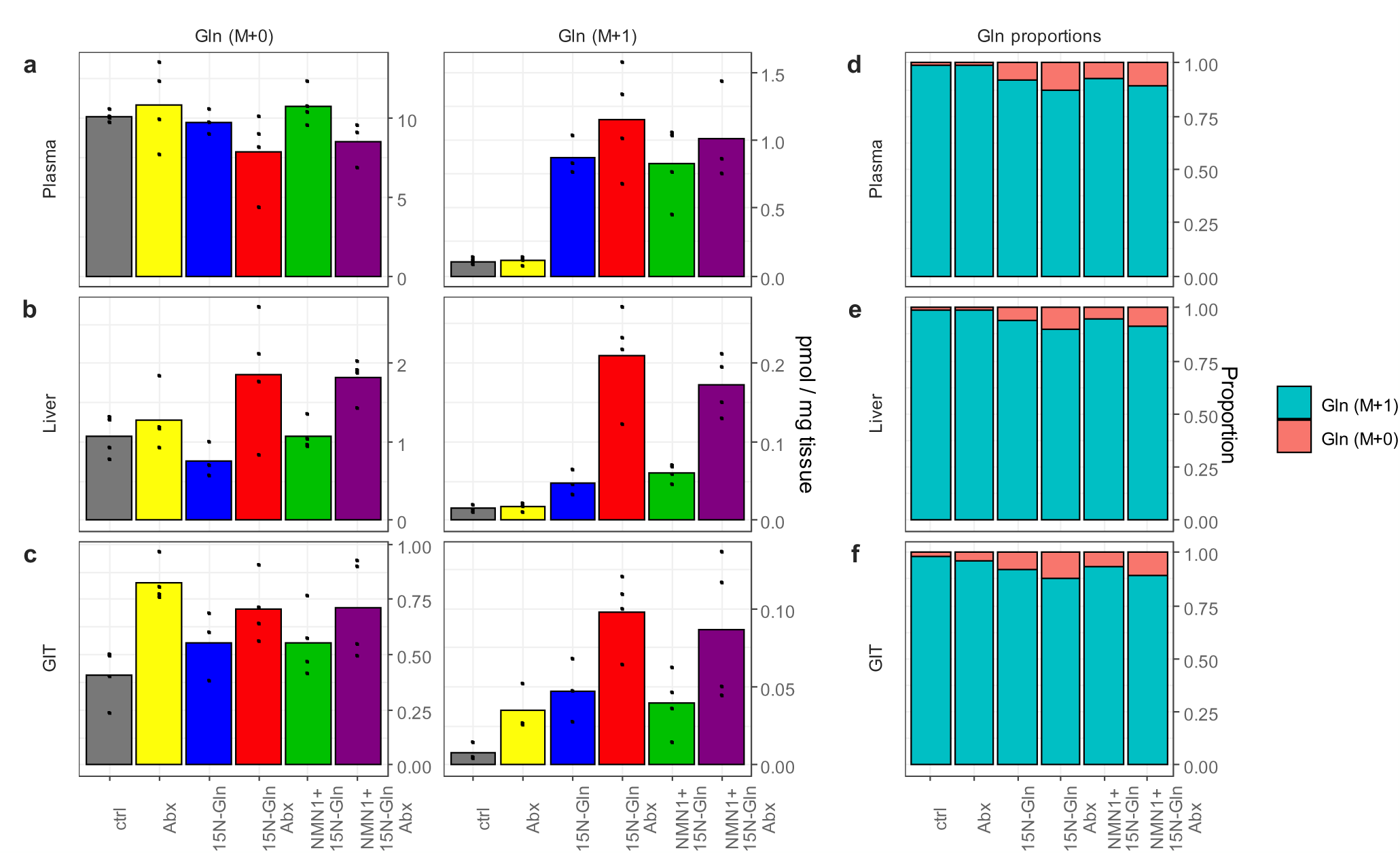
^15^N-glutamine labelling *in vivo* during NMN1 (M+6) treatment – related to main Figs. 5, 6. Labelling of the total Gln pool in (a) plasma, (b) liver and (c) GIT reached a maximum of around 10-15%, impacting the formation of M+7 labelled NAD^+^.

**Supplementary Figure 8.**
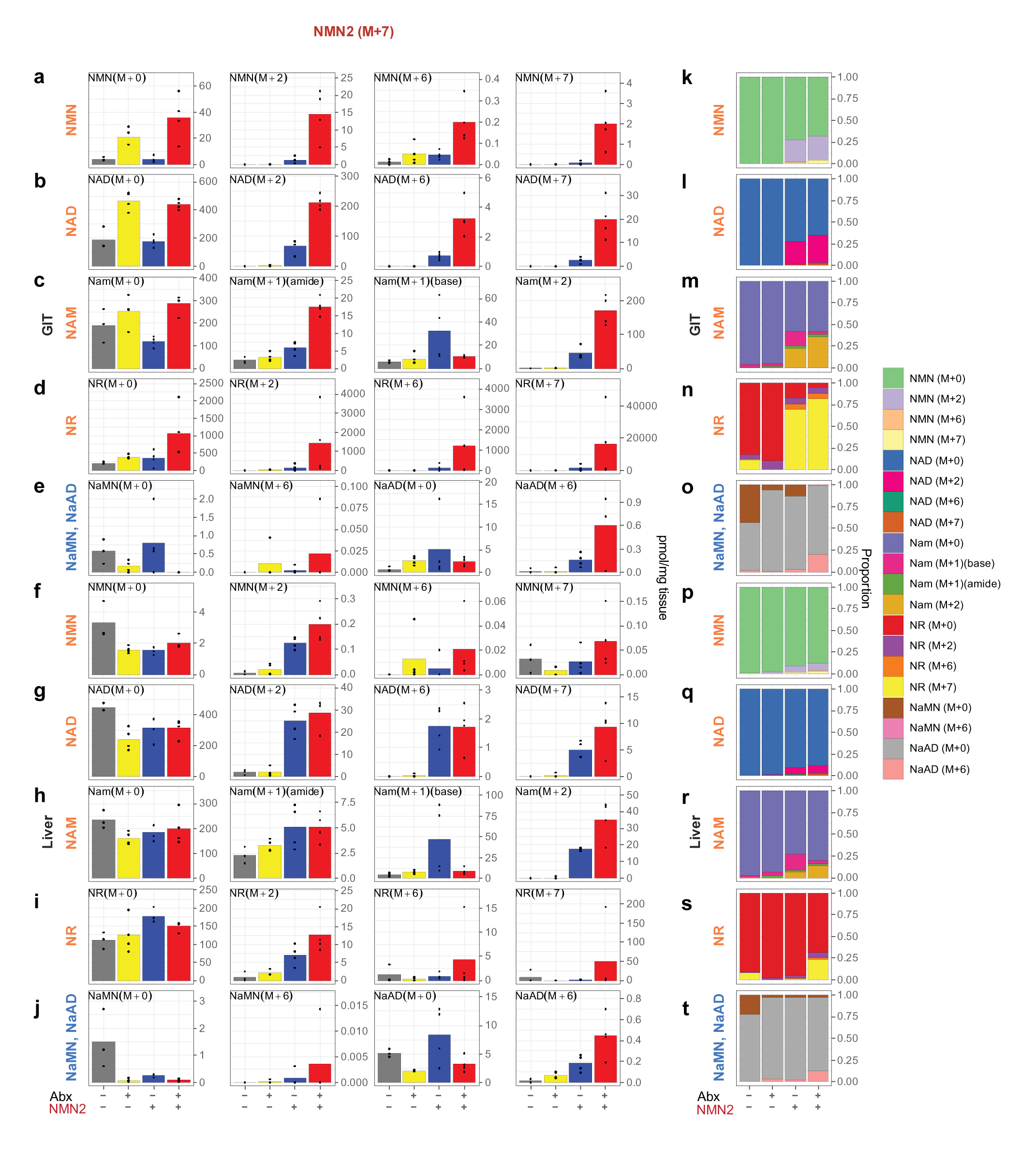
Incorporation of NMN2 (M+7) into the NAD metabolome *in vivo*. Animals were treated with antibiotics (Abx) to deplete the microbiome, followed by an oral gavage (50 mg/kg) of **NMN2** (M+7). Four hr later, GIT (a-g), liver (h-n) and plasma (o) were rapidly preserved for targeted metabolomics analysis to identify labelling of (a, h, o) Nam, (b, i) NMN, (c, j) NR, (d, k) NAD^+^, (e, l) NaMN, (f, m) NaR and (g, n) NaAD. Data presented as stacked bars of each isotopologue, with raw data points for each isotopologue overlaid on bar charts. (p, q) Relative molar abundance of all NAD metabolites in (p) GIT and (q) liver, including endogenous (M+0), intact labelled and partially labelled (“part”) isotopologues. n=3-4 animals per group.

## SUPPLEMENTARY METHODS

### Synthesis of isotope labelled NMN

The isotopes used here were generated through a two-step process starting with the custom synthesis of nicotinamide labelled with ^15^N at the nitrogen base and amide positions. This custom isotope labelled version of nicotinamide was then used with [U5]-^13^C- ribose which was ^13^C labelled at all five carbon positions (Cambridge Isotope Laboratories, cat. no. CLM-3652) and ATP in an enzyme-based protocol using recombinant phosphoribosyl synthetase (PRS) and recombinant nicotinamide phosphoribosyl transferase (NAMPT) to synthesise NMN. The two enzymes were added into the reaction buffer that contains 1 mM ribose, 1 mM nicotinamide, 3 mM ATP, 1 mM dithiothreitol, 10 mM MgCl_2_ and 50 mM Tris-HCl (pH 7.5) and incubated at 37℃ for 30 min. The reaction was terminated with the addition of 0.01% Trichloroacetic acid (TCA). The purification was proceeded with size-exclusion columns and ion exchange columns. Isotope labelled NMN samples of >95% purity were concentrated by lyophilization, and labelling confirmed by mass spectrometry (Supp. Fig. 2).

### Animal experiments

All experiments were performed according to procedures approved by UNSW Animal Care and Ethics Committee (ACEC) under ethics protocol 18/134A. The UNSW ACEC operates under the animal ethics guidelines from the National health and Medical Research Council (NHMRC) of Australia. Mice were fed standard chow *ad libitum* and housed under a 12-hr light/12-hr dark cycle in a temperature-controlled room (22 ± 1 °C) at 80% humidity in individually ventilated cages. Four-week old female C57BL/6J mice were acclimatised for one week prior to treatment and body weight matched before random assignment into groups. For antibiotic treatment, mice were administered a cocktail of antibiotics consisting of vancomycin (0.5 g/L; Sigma SBR00001), neomycin (1 g/L; Sigma N6386), ampicillin (1 g/L; Sigma A9393) and metronidazole (1 g/L; Sigma, M3761) (VNAM) with addition of sucrose (3 g/L; Bundaberg Sugar) to increase palatability for 4 days, and switched to ampicillin (1 g/L) with sucrose (3 g/L) for an additional week, which can reduce gut bacterial density by 1000-fold ^71^. During treatment with the VNAM combination there was a reduction in water consumption (below), which was the reason for the subsequent switch to ampicillin alone. Sucrose treatment (3 g/L) was maintained as a vehicle control in animals that did not receive antibiotic treatment.

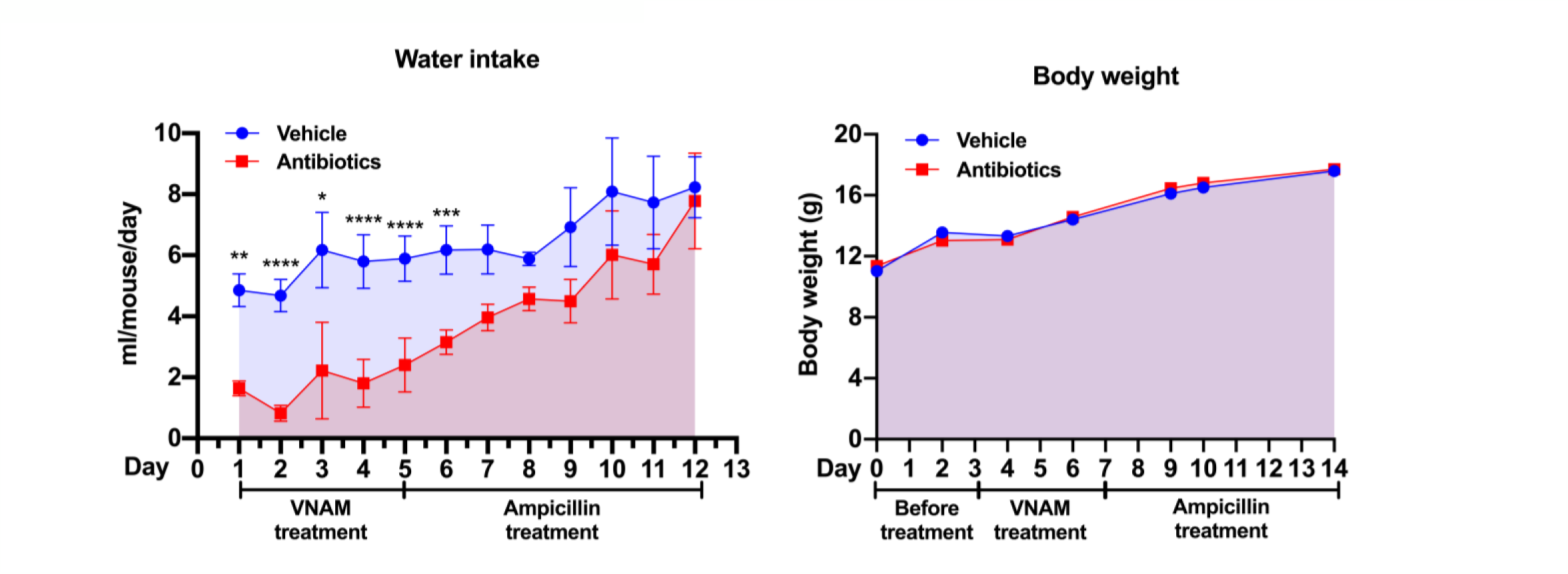

*Above: Water intake and body weight of mice during antibiotics treatment*.

To maintain consistency between the three cohorts presented here (unlabelled NMN – Fig. 1, NMN1 – Fig. 5, NMN2 – Fig. 6), this antibiotic treatment protocol was used for all *in vivo* experiments (Figs. 1, 5, 6). For NMN treatment, mice received a single oral gavage of NMN1 (Fig. 5) or NMN2 (Fig. 6) isotopes at 50 mg/kg, or for unlabelled NMN (Fig. 1) at 500 mg/kg, with water vehicle used as a control. Four hours later, animals were placed under isofluorance anaesthesia, and blood was obtained by cardiac puncture, euthanasia by cervical dislocation, rapid dissection and snap freezing of tissues. Gavages were staggered between mice in alternating treatment groups to avoid any experimental bias. On the day of cull mice were all 5-6 weeks old. Differences in NMN dosing between unlabelled and isotope labelled NMN were due to limited availability of isotope labelled NMN.

### Blood plasma collection and preparation for mass spectrometry

Approximately 1 mL of blood was collected via cardiac puncture in anaesthetised mice (1-2% isoflurane) into 1.5 mL eppendorf tubes prefilled with 10 µL of EDTA (0.5 M) and mixed thoroughly with a pipette to prevent clotting. Blood samples were then spun at 2000 *g* for 10 min and the top layer was transferred to a new tube and snap frozen immediately in liquid nitrogen. All samples were stored in -80°C until further processing. On the day of sample acquisition plasma samples were thawed on ice and 20 µL of plasma was added to 80 µL of extraction buffer (acetonitrile:methanol) with an internal standard mixture containing MES, CSA and thymine-d4. Samples were vortexed and centrifuged at 16,000*g* for 10 mins at 4°C and the supernatant was transferred to a new eppendorf tube and dried down completely using a speed vacuum concentrator (Savant SpeedVac ® SPD140DDA, Thermo Scientific). The resulting pellet was then resuspended in 30 µL of LC-MS-grade water and centrifuged as above and the supernatant analysed promptly by LC-MS.

### Gastrointestinal and liver tissue collection and preparation for mass spectrometry

Intestinal contents (small intestine and colon without cecum) were resected and flushed with ice cold 1 x phosphate buffered saline (PBS) to clear faecal contents before snap freezing immediately with liquid nitrogen. Livers were resected and weighed before being rinsed in ice cold 1x PBS and snap frozen in liquid nitrogen. All tissue samples were stored in -80 °C until further processing. Frozen tissue samples were crushed using a mortar and pestle on liquid nitrogen and approximately 50 mg was weighed into tubes containing ceramic beads (Precellys, Bertin Technologies, France) to which 500 µl of cold (-30 °C) extraction buffer (acetonitrile:methanol:water, 2:2:1) with internal standard mixture as above was added. All samples were homogenised using an automated tissue homogeniser (Precellys24, Bertin Technologies, France) at 5,000-6,000 rpm for 15 seconds and immediately centrifuged at 16,000*g* for 10 mins at 4 °C. The supernatant was transferred to a new tube and dried down completely using a speed vacuum concentrator (Savant SpeedVac ® SPD140DDA, Thermo Scientific). All samples were resuspended in 50 µL of LC-MS-grade water, centrifuged as above and the supernatant analysed promptly by LC-MS.

### Bacterial culture and NMN treatment

A stab culture of the *E. coli* strain OP50 was inoculated into sterile Luria-Bertani (LB) broth (10 g/L tryptone, 5 g/L yeast and 10 g/L sodium chloride in deionized water) under aseptic conditions and incubated overnight at 37℃ on a shaking platform set at 200 rpm. To measure the growth rate of *E. coli*, the overnight culture was sub-cultured (1:200) into sterile LB broth in a new flask and the optical density was measured at 600 nm (OD_600_) every 20 minutes (approximate doubling time) and samples were collected during the early-mid exponential growth phase (OD600 < 0.70), as bacterial enzymes are more active during exponential growth phase than stationary phase ^72^. For samples, the overnight culture was sub-cultured (1:200) and aliquoted into smaller volumes. The cultures were then supplemented with either vehicle (water) or M+6 labelled NMN (0.1 mM) and OD600 measured at time zero (before NMN), time zero (after NMN), and 140, 160 and 180 minutes after supplementation with NMN. The supernatant of cells was separated from the cells via centrifugation (5000 *g* for 10 minutes at 4℃) and stored immediately at -30℃. Meanwhile, the cell pellet was resuspended in cold (4°C) saline solution (0.9% NaCl) and centrifuged as above, to rinse away residual media before storage at -30℃. The OD_600_ was measured for each sample and used to normalise metabolite levels after LC-MS/MS analysis.

### Primary hepatocyte culture

Primary hepatocytes were obtained as described previously ^73^. Male Sprague Dawley rats (250 grams, Animal Resources Centre, Perth, WA, Australia) were maintained on a 12:12 h day-night cycle, with water and food supplied *ad libitum*. Under deep non-recoverable general anaesthesia (75 mg/kg ketamine, 10 mg/kg xylazine, intraperitoneal administration) rats underwent laparotomy. The portal vein was cannulated *in situ* and the liver perfused initially with carbogen-saturated perfusion media (final: NaCl 138 mM, HEPES 25 mM, D-glucose 5.6 mM, KCl 5.4 mM, Na_2_HPO_4,_ 0.34 mM, KH_2_PO_4_ 0.44 mM, NaHCO_3_ 4.17 mM, EDTA 0.5 mM, pH 7.4, 37°C, 25 ml/min flow rate). The inferior vena cava was cut to allow efflux. After 4 mins, the carbogen-saturated perfusion media was changed to the collagenase containing buffer (final: NaCl 138 mM, HEPES 25 mM, D-glucose 5.6 mM, KCl 5.4 mM, Na_2_HPO_4,_ 0.34 mM, KH_2_PO_4_ 0.44 mM, NaHCO_3_ 4.17 mM, CaCl_2_ 2 mM, collagenase II (Sigma, 15950-017), pH 7.4, 37°C, 25 ml/min flow rate) for 6 mins. The inferior vena cava was clamped at least 10 times during the collagenase digestion (preventing efflux) resulting in liver swelling that allows a better digestion.

Following the collagenase digestion, the liver was removed and place on ice in 20 ml Williams’ Medium E (Life-technologies, Waltham, MA, USA). The hepatocytes were gently dispersed in the medium and the cells filtered through a 100 µm cell strainer. Hepatocytes were washed and diluted Williams’ Medium E and plated (6-well plates) at 10^6^/2ml/well. After 4 hrs of incubation the culture medium was changed to MOPS buffer (final, NaCl 128 mM, MOPS 23.9 mM, KCl 6 mM, MgSO_4_.7H_2_O 1.18 mM, CalCl_2_ 1.29 mM, glucose 5 mM, BSA (FFA) 0.2 %, pH 7.4) and cells incubated overnight. Following overnight incubation, cells were incubated in M199 media without glutamine (Sigma M2154), supplemented with either unlabelled (Sigma) or ^15^N-amide labelled glutamine (Cambridge Isotope Laboratories NLM-557) at 4 mM, in the presence or absence of NMN1 or NMN2 isotopes at 200 µM for 24 hr, following which samples were preserved for metabolomic analysis (Fig. 3).

### Preparation of NAD^+^ metabolite standards

NAD^+^ metabolites were serially diluted starting from a concentration of 100 µM to 0.39 µM. The same volume (500 µL) of extraction buffer (acetonitrile:methanol:water) was added and vortexed before centrifuging and transferring to new tube ready to be dried down as above. The subsequent steps were the same as preparing the tissue samples as above. All standards and samples were processed on the same day to reduce any experimental bias or variability. Standard curves used to calculate absolute concentrations are shown on the following page (Methods Fig. 2) and are available in supplementary raw data files.

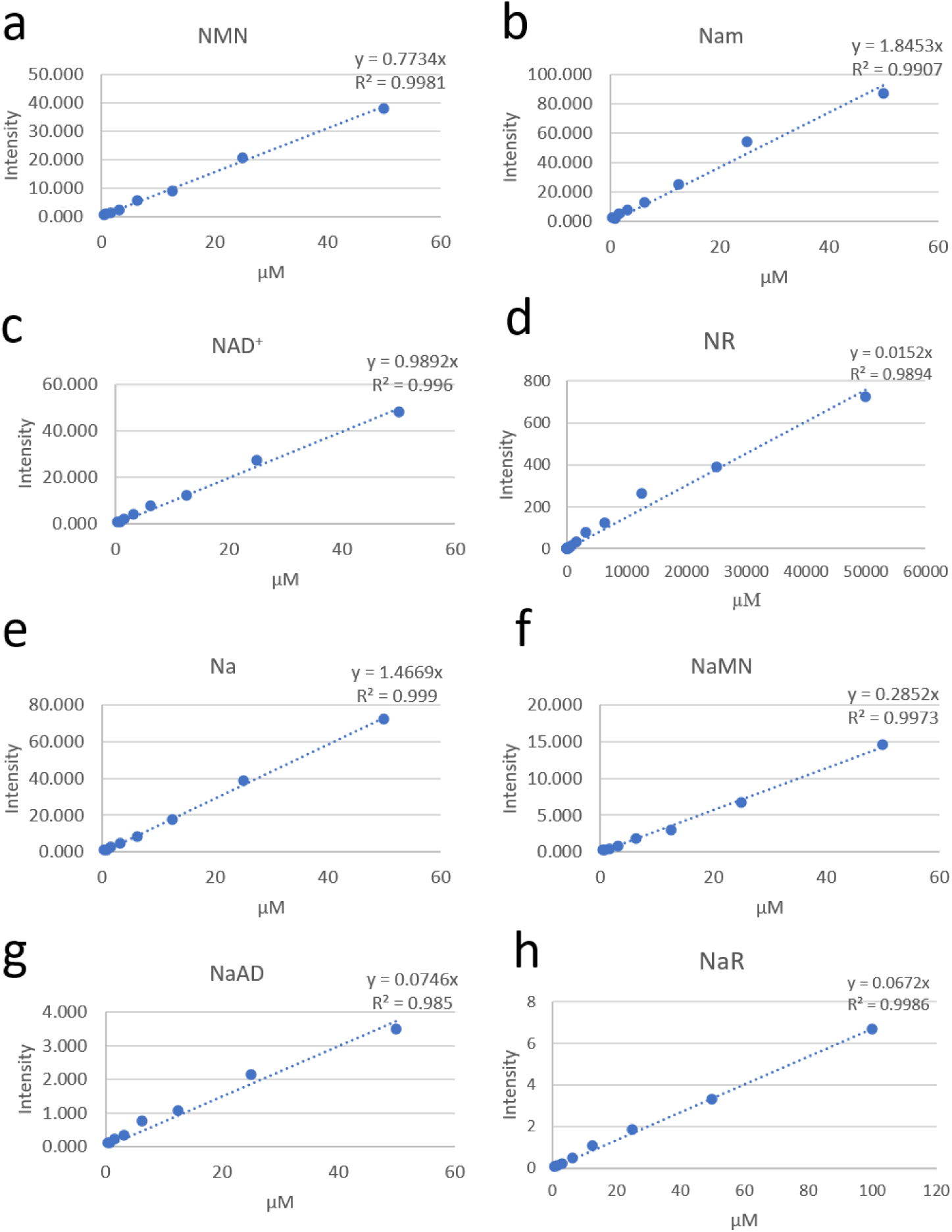

*Above: NAD^+^ metabolite standard curves. (a) NMN (b) Nam (c) NAD^+^ (d) NR (e) Na (f) NaMN (g) NaAD (h) NaR. Standard curves were serially diluted from 50 µM to 0.39 µM. Shown here from 50 µM to 0.39 µM for all metabolites except for NR which is shown from 50 mM to 0.39 µM, due to the unexpectedly high concentrations of NR in the GIT (Fig. 1, 5, 6) and NaR from 100 µM to 0.39 µM*.

### Mass spectrometry

The LC-MS method was performed using 1260 Infinity LC System (Agilent) coupled to QTRAP 5500 (AB Sciex) mass spectrometer. LC separation by gradient elution was accomplished on an XBridge BEH amide column (100 mm x 2.1 mm, 3.5 µm particle size, Waters Corporation) at room temperature. For the mobile phase, Solvent A is 95%:5% H_2_O:acetonitrile containing 20 mM ammonium acetate and 20 mM acetic acid, and solvent B is acetonitrile. The flow rate was 200 µL/min, with the percentage of solvent B set at 85% (0 min), 85% (0.1 min), 70% (10 min), 15% (13 min), 15% (17 min), 85% (17.5 min), 85% (30 min) (Supplementary Table 1). Injection volume was 2.5 µL. Ion source was set at 350 °C and 4500 V with polarity switching. Mass isotopologues of metabolites were acquired by MS^2^, using the unscheduled multiple reaction monitoring (MRM) mode with a dwell time of 40 ms. The MS parameters (declustering potential, collision energy and cell exit potential) (Supplementary Table 2a) and MRM transitions were calibrated based on the monoisotopic mass of chemical standards (Supplementary Figure 11). Data processing was performed using MSConvert (version 3.0.18165-fd93202f5) and in-house MATLAB scripts. Deconvolution scripts were developed to resolve overlapping NAAD-NAD, NAR-NR and NAMN-NMN peaks using MATLAB’s Optimisation Toolbox. Representative chromatograms are shown below.

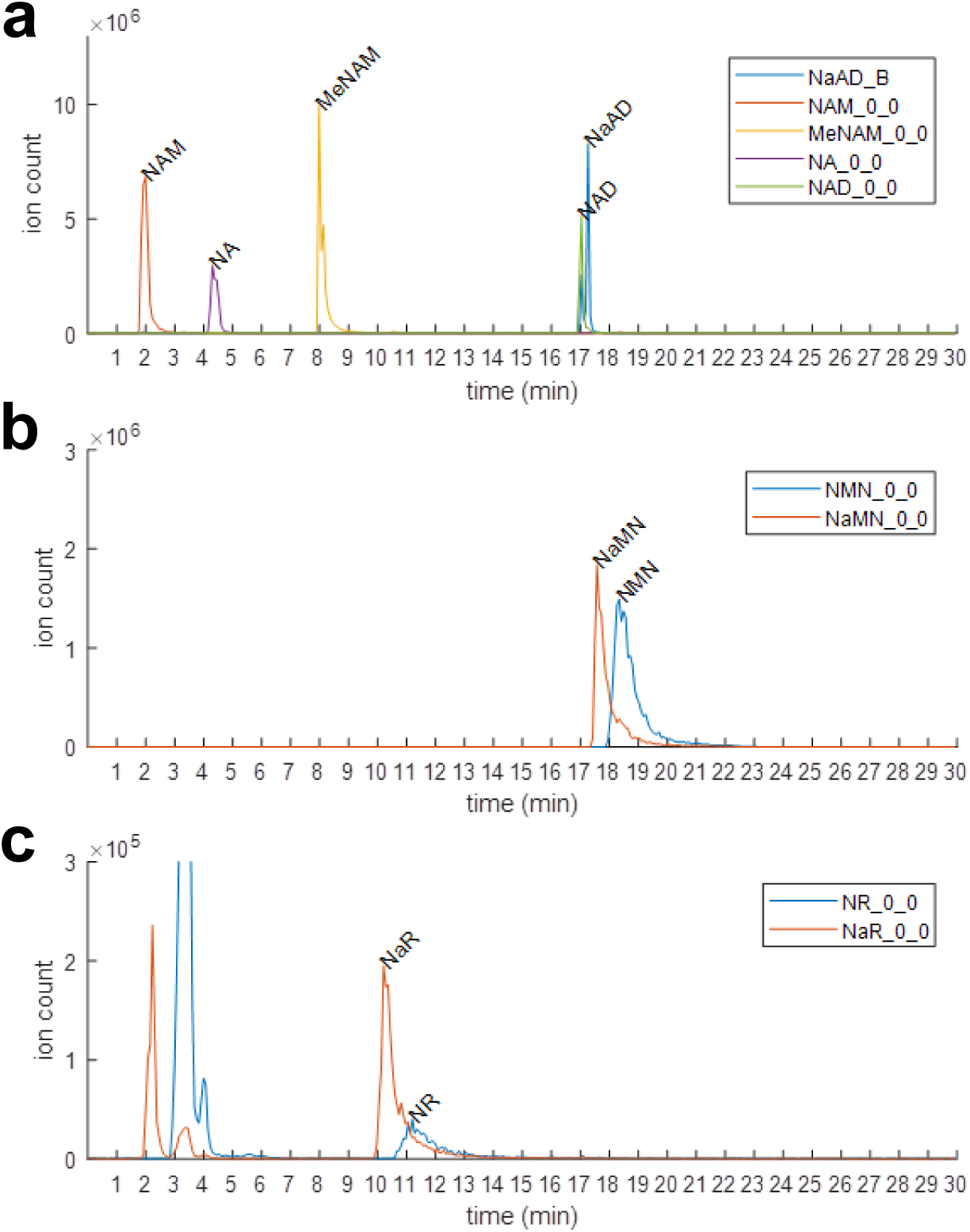

*Left: chromatograms represent individual peaks (ion count) from 100µM standard solutions at each respective retention times for **(a)** NAM, NA, MeNAM, NAD^+^ and NAAD, **(b)** NAMN and NMN, **(c)** NAR and NR*.

### Isotope correction

Natural isotope abundance correction was performed using an in-house MATLAB script based on *IsoCorrectoR* but modified for our data^74^. Our tandem MS/MS data (i.e., MRMs) targeted the most informative fragments where label(s) reside, as such is not a complete mass distribution vector. The correction procedure needs to subtract enrichments of natural atoms from our "unnatural isotope- centric MRM". Briefly, we first resolved molecular identities of the product ion and the neutral loss(es) for each metabolite, since the ^13^C and ^15^N labels can reside in either. Secondly, we used a least-square procedure to simulate our MRM data vector using mass distributions of a given product ion and neutral loss species by optimising the label(s) present in both. The result from the best fit gave the corrected mass distribution vectors of both the product ion and neutral loss, which in turn were used to calculate the corrected MRMs matching what was acquired.

### Statistical analysis for mass spectrometry

All data are presented as mean ± standard deviation (s.d.), except for estimation plots shown in Fig. 5, which utilise 95% confidence intervals. Statistical significance for the 2x2 design in Fig. 1 was performed using a two-way ANOVA with a Sidak’s multiple comparisons test to determine differences between samples. Data in Fig. 5 were analysed by permutational t-test between NMN isotope treated groups due to the absence of detection of labelled metabolites in animals that did not receive NMN isotopes. Statistics were performed on GraphPad Prism software (version 8.2.1) and using the *MKinfer* package for permutational t-tests in R. P values less than 0.05 were considered statistically significant. All data analyses are available as an .xml file and an annotated R script available on our Mendeley data site. For *in vivo* experiments (Fig. 1, 5-8, Supp. Fig. 4-7), each data point represents tissues from a separate animal, while each data point for *in vitro* experiments (Fig. 2, 3) represents an independent biological replicate.

### DNA extraction from faecal pellets

Solid faecal pellets taken from the colonic and rectal region of the gastrointestinal tract were stored in -80℃ until further processing. DNA was extracted from frozen faecal pellets using the QIAamp® PowerFecal® DNA kit (Qiagen, Cat. No. 12830-50) according to the manufacturer’s protocol. DNA concentration was determined using the NanoDrop™ (DeNovix®, DS-11 FX) and the purity of double-stranded DNA (dsDNA) was also determined by measuring the 260/280 ratio. All DNA extracts were stored at -80℃ until further processing by 16S rRNA sequencing. Bacterial DNA was quantified (Fig 1b) by qRT-PCR of genomic DNA for bacterial 16S and mouse Rsp18, with data expressed as a ratio of the two.

### 16S Sequencing

Full length 16S rRNA genes were amplified by PCR using the Oxford Nanopore 16S Barcoding Kit (SQK-RAB204; Oxford Nanopore Technologies, Oxford, UK). Briefly, 10 ng genomic DNA, 1 µL 16S Barcode (10 µM) and 25 µL LongAmp Taq 2X Master Mix (New England Biolabs, Ipswich, MA, USA) were combined in a 50 µL reaction for PCR on a Bio-Rad T100^TM^ Thermal Cycler (Bio-Rad Laboratories Pty Ltd, Hercules, CA, USA). PCR cycling condition were as follows; initial denaturation at 95 ℃ for 1 minute, 25 cycles of denaturation at 95 ℃ for 20 seconds, annealing at 55 ℃C for 30 seconds and extension at 65 ℃ for 2 minutes before a final extension at 65 ℃ for 5 minutes. PCR products were purified as per Oxford Nanopore Technologies (ONT) protocol using AMPure XP magnetic beads (Beckman Coulter, Indianapolis, IN) and DNA quantified using the NanoDrop™ (DeNovix®, DS-11 FX). Barcodes were pooled to a total of 100 fmol in 10 μL of 10 mM Tris-HCl, pH 8.0 with 50 mM NaCl for library loading. Sequencing was performed using R9.4.1 ONT Flow Cells on the MinION^TM^ sequencing platform and data acquired using MinKNOW software version 19.10.1 (Oxford Nanopore Technologies).

### Data Analysis for 16S sequencing

Full length 16S sequencing reads acquired from MinION runs (i.e. FAST5 data) were base-called to fastq files using *Guppy* software version 3.4.4 (Oxford Nanopore Technologies). Fastq files were demultiplexed using *Porechop* (https://github.com/rrwick/Porechop) and trimmed to 1400bp with *Trimmomatic* version 0.39 ^75^. Reads were imported to *QIIME2* for dereplication and chimeric reads screened and filtered from the dataset. Operational taxonomic unit clustering was completed within *QIIME2* version 2019.7.0 ^76^ at 85% similarity to account for typical sequencing errors obtained from long-read sequencing. Taxonomy was assigned to reads using a pre-trained classifier on the SILVA 132 16S rRNA representative sequences. Data was imported into *R* version 3.6.1 with *qiime2R* version 0.99.13 (https://github.com/jbisanz/qiime2R) for visualisation and alpha diversity analysis using raw and rarefied data with the *phyloseq* version 1.30.0 ^77^ package. Scripts for command line processing and analysis in R available in Supplementary Materials. Sequencing data has been deposited in the NCBI database Sequence Read Archive (SRA) under accession numbers PRJNA635359.

## Figures

Labelling schemes and illustrator were created using BioRender.com and Adobe Illustrator. Data were prepared into figures using both GraphPad Prism and using the R packages *ggplot2, ggtext, patchwork, ggh4x* and *ggpubr*. Annotated R scripts for data presentation have been uploaded to our Mendeley site.

## SUPPLEMENTARY TABLES

**Supplementary Table 1.** Liquid chromatography (LC) separation gradient Buffer A: 95:5 (v/v) HPLC H_2_O:Acetonitrile (CH_3_CN) with 20 mM ammonium acetate (NH_4_OAc) + 20mM acetic acid (CH_3_COOH), pH 5. Buffer B: 100% Acetonitrile (CH_3_CN).

| Time (mins) | Flow rate (µl/min) | Buffer A (%) | Buffer B (%) |
| --- | --- | --- | --- |
| 0 | 200 | 15 | 85 |
| 0.1 | 200 | 15 | 85 |
| 10 | 200 | 30 | 70 |
| 13 | 200 | 85 | 30 |
| 17 | 200 | 85 | 30 |
| 17.5 | 200 | 15 | 85 |
| 30 | 200 | 15 | 85 |

**Supplementary Table 2.** MRM transitions and MS parameters of NAD^+^ metabolites and MRM internal standards (Thymidine d4, CSA, MES). Q1: parent ion, Q3: fragment ion, MRM: multiple reaction monitoring, DP: declustering potential, CE: collision energy, CXP: collision cell exit potential.

| **Metabolite_Q1_Q3** | **Q1**<br>**(m/z)** | **Q3**<br>**(m/z)** | **DP**<br>**(V)** | **CE**<br>**(V)** | **CXP**<br>**(V)** | **RT**<br>**(mins)** |
| --- | --- | --- | --- | --- | --- | --- |
| glutamine_0_0 | 147 | 44 | 51 | 73 | 10 | 13 |
| glutamine_1_0 | 148 | 44 | 51 | 73 | 10 | 13 |
| glutamine_1_1 | 148 | 45 | 51 | 73 | 10 | 13 |
| glutamine_2_1 | 149 | 45 | 51 | 73 | 10 | 13 |
| NA_0_0 | 124 | 78 | 70 | 25 | 10 | 4 |
| NA_1_1 | 125 | 79 | 70 | 25 | 10 | 4 |
| NaAD_0_0 | 665 | 428 | 139 | 35 | 38 | 17 |
| NaAD_5_0 | 670 | 428 | 139 | 35 | 38 | 17 |
| NaAD_6_0 | 671 | 428 | 139 | 35 | 38 | 17 |
| NaAD_1_0 | 666 | 428 | 139 | 35 | 38 | 17 |
| NaAD_5_5 | 670 | 433 | 139 | 35 | 38 | 17 |
| NaAD_10_5 | 675 | 433 | 139 | 35 | 38 | 17 |
| NaAD_11_5 | 676 | 433 | 139 | 35 | 38 | 17 |
| NaAD_6_5 | 671 | 433 | 139 | 35 | 38 | 17 |
| NaAD_6_6 | 671 | 434 | 139 | 35 | 38 | 17 |
| NaAD_11_6 | 676 | 434 | 139 | 35 | 38 | 17 |
| NaAD_12_6 | 677 | 434 | 139 | 35 | 38 | 17 |
| NaAD_7_6 | 672 | 434 | 139 | 35 | 38 | 17 |
| NaAD_1_1 | 666 | 429 | 139 | 35 | 38 | 17 |
| NaAD_6_1 | 671 | 429 | 139 | 35 | 38 | 17 |
| NaAD_7_1 | 672 | 429 | 139 | 35 | 38 | 17 |
| NaAD_2_1 | 667 | 429 | 139 | 35 | 38 | 17 |
| NaAD_7_7 | 672 | 435 | 139 | 35 | 38 | 17 |
| NAD_0_0 | 664 | 428 | 33 | 36 | 31 | 17 |
| NAD_5_0 | 669 | 428 | 33 | 36 | 31 | 17 |
| NAD_6_0 | 670 | 428 | 33 | 36 | 31 | 17 |
| NAD_7_0 | 671 | 428 | 33 | 36 | 31 | 17 |
| NAD_1_0 | 665 | 428 | 33 | 36 | 31 | 17 |
| NAD_2_0 | 666 | 428 | 33 | 36 | 31 | 17 |
| NAD_5_5 | 669 | 433 | 33 | 36 | 31 | 17 |
| NAD_6_6 | 670 | 434 | 33 | 36 | 31 | 17 |
| NAD_7_7 | 671 | 435 | 33 | 36 | 31 | 17 |
| NAD_1_1 | 665 | 429 | 33 | 36 | 31 | 17 |
| NAD_2_2 | 666 | 430 | 33 | 36 | 31 | 17 |
| NAD_10_5 | 674 | 433 | 33 | 36 | 31 | 17 |
| NAD_11_5 | 675 | 433 | 33 | 36 | 31 | 17 |
| NAD_12_5 | 676 | 433 | 33 | 36 | 31 | 17 |
| NAD_6_1 | 670 | 429 | 33 | 36 | 31 | 17 |
| NAD_7_1 | 671 | 429 | 33 | 36 | 31 | 17 |
| NAD_8_1 | 672 | 429 | 33 | 36 | 31 | 17 |
| NAD_11_6 | 675 | 434 | 33 | 36 | 31 | 17 |
| NAD_12_6 | 676 | 434 | 33 | 36 | 31 | 17 |
| NAD_13_6 | 677 | 434 | 33 | 36 | 31 | 17 |
| NAM_0_0 | 123 | 80 | 80 | 30 | 25 | 2 |
| NAM_1_1 | 124 | 81 | 80 | 30 | 25 | 2 |
| NAM_2_1 | 125 | 81 | 80 | 30 | 25 | 2 |
| NAM_1_0 | 124 | 80 | 80 | 30 | 25 | 2 |
| NaMN_0_0 | 336 | 124 | 66 | 28 | 11 | 19 |
| NaMN_5_0 | 341 | 124 | 66 | 28 | 11 | 19 |
| NaMN_6_1 | 342 | 125 | 66 | 28 | 11 | 19 |
| NaMN_1_1 | 337 | 125 | 66 | 28 | 11 | 19 |
| NaR_0_0 | 256 | 124 | 41 | 27 | 6 | 10 |
| NaR_5_0 | 261 | 124 | 41 | 27 | 6 | 10 |
| NaR_6_1 | 262 | 125 | 41 | 27 | 6 | 10 |
| NaR_1_1 | 257 | 125 | 41 | 27 | 6 | 10 |
| NMN_0_0 | 335 | 123 | 48 | 24 | 11 | 19 |
| NMN_5_0 | 340 | 123 | 48 | 24 | 11 | 19 |
| NMN_6_1 | 341 | 124 | 48 | 24 | 11 | 19 |
| NMN_7_2 | 342 | 125 | 48 | 24 | 11 | 19 |
| NMN_1_1 | 336 | 124 | 48 | 24 | 11 | 19 |
| NMN_2_2 | 337 | 125 | 48 | 24 | 11 | 19 |
| NR_0_0 | 255 | 123 | 64 | 30 | 13 | 11 |
| NR_5_0 | 260 | 123 | 64 | 30 | 13 | 11 |
| NR_6_1 | 261 | 124 | 64 | 30 | 13 | 11 |
| NR_7_2 | 262 | 125 | 64 | 30 | 13 | 11 |
| NR_1_1 | 256 | 124 | 64 | 30 | 13 | 11 |
| NR_2_2 | 257 | 125 | 64 | 30 | 13 | 11 |
| Thymine-d4 | 129 | 42 | -115 | -52 | -11 | 2 |
| CSA | 231 | 80 | -170 | -40 | -13 | 2 |
| MES | 196 | 100 | 140 | 31 | 25 | 4 |

